# Syk activation during FcγR-mediated phagocytosis involves both Syk palmitoylation and desulfenylation

**DOI:** 10.1101/2025.05.26.656092

**Authors:** Maxime Jansen, Jean-Marc Strub, Laurent Chaloin, Peter Coopman, Bruno Beaumelle

## Abstract

The non-receptor Spleen tyrosine kinase Syk acts downstream of several receptors of the immune system such as the FcγR. Syk is composed of a kinase domain and two SH2 domains that interact with the bi-phosphorylated ITAMs motifs of the FcγR upon phagocytosis. This results in the activation of Syk by auto- phosphorylation, triggering phosphorylation of several downstream targets in a process that will culminate in F-actin polymerization and phagocytosis of the IgG-opsonized target.

We found that Syk is S-acylated upon phagocytosis by macrophages. Palmitoylation is performed on a single Syk-Cys by the protein S-acyl transferase DHHC5 that specifically associates with Syk upon phagocytosis. Syk palmitoylation is required for Syk localization to the phagocytic cup, Syk phosphorylation/activation, Cdc42 recruitment to the cup, F-actin polymerization and phagocytosis.

We also observed that another Syk-Cys residue is modified by sulfenylation. Mutation of the sulfenylated Cys that belongs to a redox-motif inactivated the Syk catalytic activity and phagocytosis. We found that Syk desulfenylation occurs during phagocytosis. Molecular dynamics studies indicated that desulfenylation increased the mobility and exposure of a loop within the Syk interdomain B, likely facilitating phosphorylation of key Syk-Tyr residues by upstream effectors such as Src kinases.

We thus propose an original updated model for Syk activation during FcγR-mediated phagocytosis that involves both Syk palmitoylation and desulfenylation.

## Introduction

The Syk tyrosine kinase acts downstream of several receptors that contain in their cytoplasmic tail an immunoreceptor tyrosine-based activation motif (ITAM), or a hemITAM that are short sequences with two or one phosphorylable Tyr residues, respectively. The ITAM can also be present on a receptor-associated protein. These receptors include the Fc receptor (FcR), the T-cell and B-cell antigen receptors (TCR and BCR), C-type lectins (CLEC), and the microglial triggering receptor expressed on myeloid cells 2 (TREM2). Syk plays a key role in adaptive immune receptor signaling, but is also involved in other cellular functions such as adhesion, osteoclast maturation, fungal pathogens and *Mycobacterium tuberculosis* recognition, necrosis and clearance of amyloid beta by microglia. Syk also plays a role in autoimmune diseases and haematological cancers (Ennerfelt *et al*, 2022; Mocsai *et al*, 2010; Singaram *et al*, 2023).

Phagocytosis by the FcγR has often been used to study the mechanism of Syk activation. Binding of IgG-opsonized particles triggers receptor activation and Src family kinases then phosphorylate the Tyr residues within ITAMs. Syk is a 72 kDa non-receptor protein that contains a kinase domain and two Src homology 2 (SH2) domains that maintain the kinase domain in an autoinhibitory conformation. Binding of the Syk SH2-domains to bi-phosphorylated ITAMs results in Syk activation and phosphorylation of Tyr residues, with ten of them being autophosphorylated. Binding of the second SH2 domain to phosphatidylinositol (3,4,5) triphosphate (PIP3) is also important for Syk activation and function (Singaram *et al*., 2023). Syk activation takes place rapidly after phagocytosis onset, peaking after ∼5 min (Raeder *et al*, 1999). Syk then orchestrates downstream activation pathways involving Vav family members, PLCγ and phosphoinositide 3-kinases (PI3Ks) thereby enabling Rac1 and Cdc42 activation and actin polymerization leading to pseudopod extension and particle engulfment, enabling successful phagocytosis (Freeman & Grinstein, 2014; Mylvaganam *et al*, 2021). Unlike Src family kinases, Syk is strictly necessary for phagocytosis, probably because Syk can to some extend phosphorylate ITAMs (Mocsai *et al*., 2010).

The current Syk activation model involves conformational changes linked to phosphorylated-ITAMs and PIP3 binding and Syk phosphorylation. These changes essentially affect the interdomain-B region of Syk (Bradshaw *et al*, 2024; Mansueto *et al*, 2019; Singaram *et al*., 2023).

Protein S-acylation (also termed palmitoylation) is a post-translational modification of cysteine residues involving the modification of the Cys-SH group by a fatty acid, most often palmitate, that becomes attached by a thioester bond to the protein. This modification is catalyzed by S-palmitoyl acyl transferases. These membrane-embedded enzymes contain a DHHC catalytic site and are often called DHHCs. In humans, among the 23 DHHCs, only 2-3 of them (depending on cell types) localize to the plasma membrane (Chopard *et al*, 2018). Palmitoylation can be reversed by acyl protein esterase (APT1 or APT2) (Won *et al*, 2018), enabling the cell to use palmitoylation to reversibly attach proteins to membranes. A number of proteins involved in phagocytosis are S-acylated. This is the case for Src family kinases and Rac1 (Dixon *et al*, 2021).

The activity of Src kinases is also regulated by the sulfenylation of two cysteine residues that results in the formation of Cys-SOH. These sulfenylation events were found to modulate the conformation of the Src protein (Heppner *et al*, 2018). Zap70, the second member of the Syk tyrosine kinase family, is expressed in T-cells and NK cells and shares ∼54% of homology with Syk (Turner *et al*, 2000). Zap70 was found to be sulfenylated and this modification was found to modulate both the function and stability of the protein (Thurm *et al*, 2017). Zap70 was also reported to be palmitoylated (Tewari *et al*, 2021), but this observation could not be confirmed in a later study (Schultz *et al*, 2022). Our aim was to explore the eventual palmitoylation and sulfenylation of the Syk kinase, and their potential involvement in Syk activation during phagocytosis.

Here we show that Syk is desulfenylated and S-acylated upon phagocytosis. We identified the modified cysteine residues and the DHHC enzyme responsible for Syk palmitoylation. Both desulfenylation and palmitoylation are required for Syk to enable phagocytosis.

## Results

### Identification of cysteine residues required for Syk to enable phagocytosis

To examine the role of Syk Cys residues in phagocytosis we first generated a ΔSyk RAW 264.7 mouse cell line using the CRISPR/Cas9 technique (Figure 1A). This ΔSyk cell line, termed ΔSyk RAW cells could not phagocytose IgG-opsonized latex beads anymore (Figure 1B and Figure S1), confirming the key role of Syk in phagocytosis (Raeder *et al*., 1999). This phagocytosis defect could be restored by transient transfection with a vector expressing EGFP-mSyk WT or the Cys to Ser mutants of the first seven Cys residues of mSyk. Nevertheless, mutation of the two last Cys residues induced a 70-90% drop in phagocytosis efficiency compared to WT mSyk (Figure 1B). These two mutants were poorly expressed compared to the other Cys to Ser mutants, indicating that these Cys are required for Syk stability (Figure S2). Throughout this study we adapted protocols so that this weak expression would not affect experimental results. As negative controls for phagocytosis assays we used a catalytically-dead version of mSyk (K396R, (Richards *et al*, 1996)) and a mutant in which both SH2 domains were inactivated (R41A-R194A, (Qin *et al*, 1998)). They were unable to support phagocytosis. Mutation of mSyk-PIP3 binding site R219A-K221A, (Singaram *et al*., 2023), termed RK/AA, inhibited phagocytosis efficiency by ∼ 60 %.

**Figure 1.**
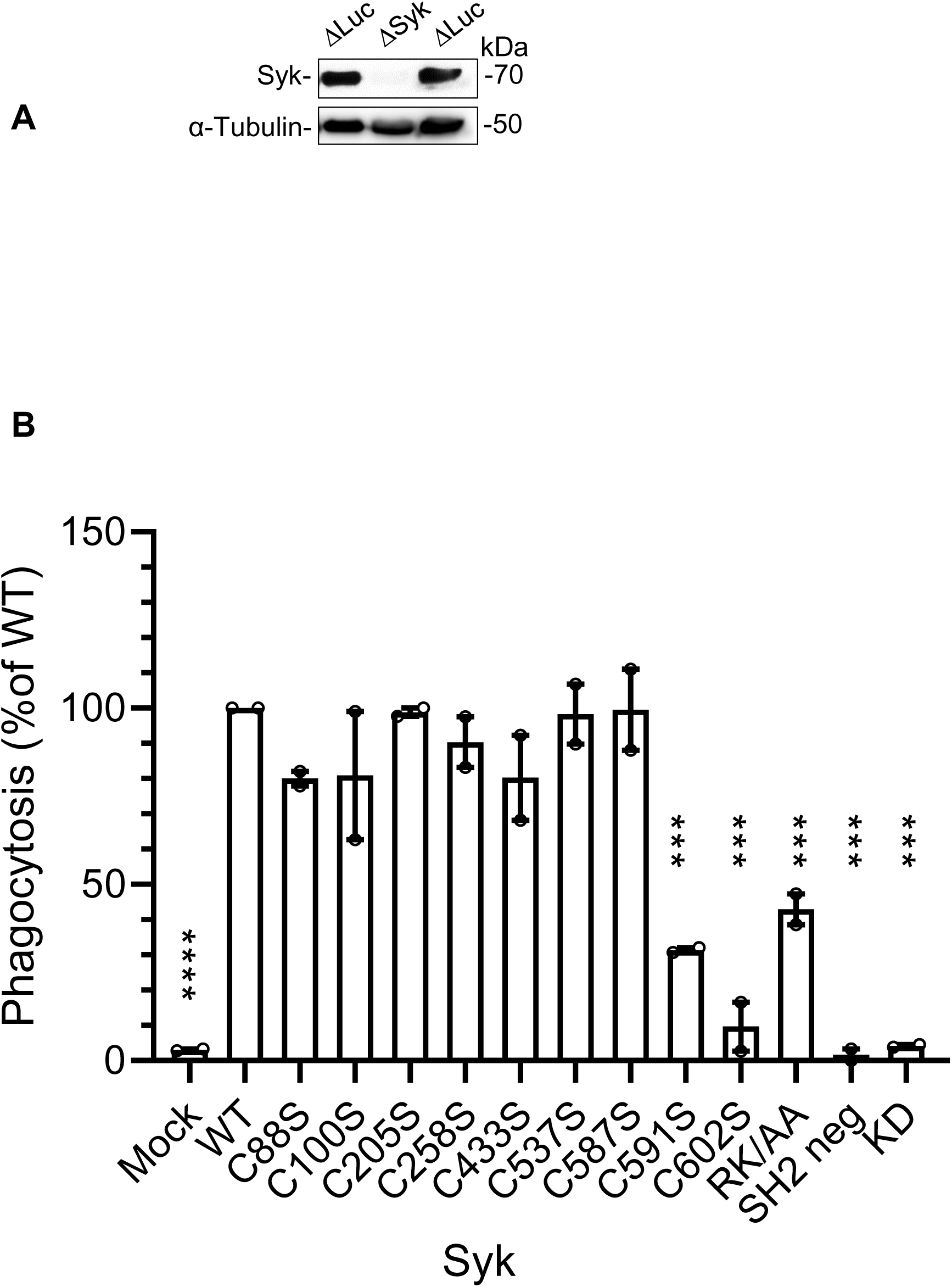
Role of Syk-Cys in phagocytosis. **A**, the ΔSyk Raw cell line does not express Syk. Western blot of cells following Syk or Firefly luciferase knockout in RAW cells using CRISPR/Cas9. **B**, Phagocytosis efficiency of Cys to Ser mutants of mSyk following transfection of EGFP-mSyk in ΔSyk RAW cells. Cells were allowed to phagocytose IgG-opsonized 3µm-diameter latex beads for 15 min at 37°C, before counting beads inside and outside cells. See Figure S1 for details.

### Syk is palmitoylated on its penultimate cysteine residue

Based on the weak biological activity (Figure 1B) and expression level of the corresponding C591S and C602S mutants (Figure S2), we wondered what could be the role of these two last Cys residues in the Syk biological activity. The CSS-palm prediction software (Ren *et al*, 2008) suggested that mSyk could be palmitoylated on Cys 591. To examine whether Syk could be palmitoylated we first used human monocyte-derived primary macrophages (hMDMs) and the acyl-biotin exchange (ABE) technique (Chopard *et al*., 2018). This technique is based on the selective cleavage of the thioester bond by hydroxylamine before attaching a biotin to the unmasked SH group, enabling to purify on streptavidin-agarose the previously palmitoylated proteins that can be identified by Western blot (Wan *et al*, 2007). We monitored Syk palmitoylation in resting macrophages or macrophages that were allowed to phagocytose IgG-opsonized targets (Sheep Red Blood Cells, SRBCs) for 5 min, which is the time that allows maximum Syk activation (Raeder *et al*., 1999). We observed that Syk is palmitoylated only upon phagocytosis (Figure 2A). Similar data were obtained using ΔSyk RAW macrophages transfected with WT mSyk (Figure 2B). When the Cys mutants were used we observed that the mutant of the last Syk-Cys (C602S) was also palmitoylated upon phagocytosis. The C591S mutant showed significant palmitoylation in the absence of phagocytosis but this level was not increased upon phagocytosis, indicating that mouse Syk is palmitoylated on C591. Such an artefactual protein palmitoylation upon mutation of the palmitoylation site has been previously observed (Abrami *et al*, 2006). The RK/AA mutant unable to bind PIP3 was palmitoylated indicating that PIP3 binding is not required for palmitoylation.

**Figure 2.**
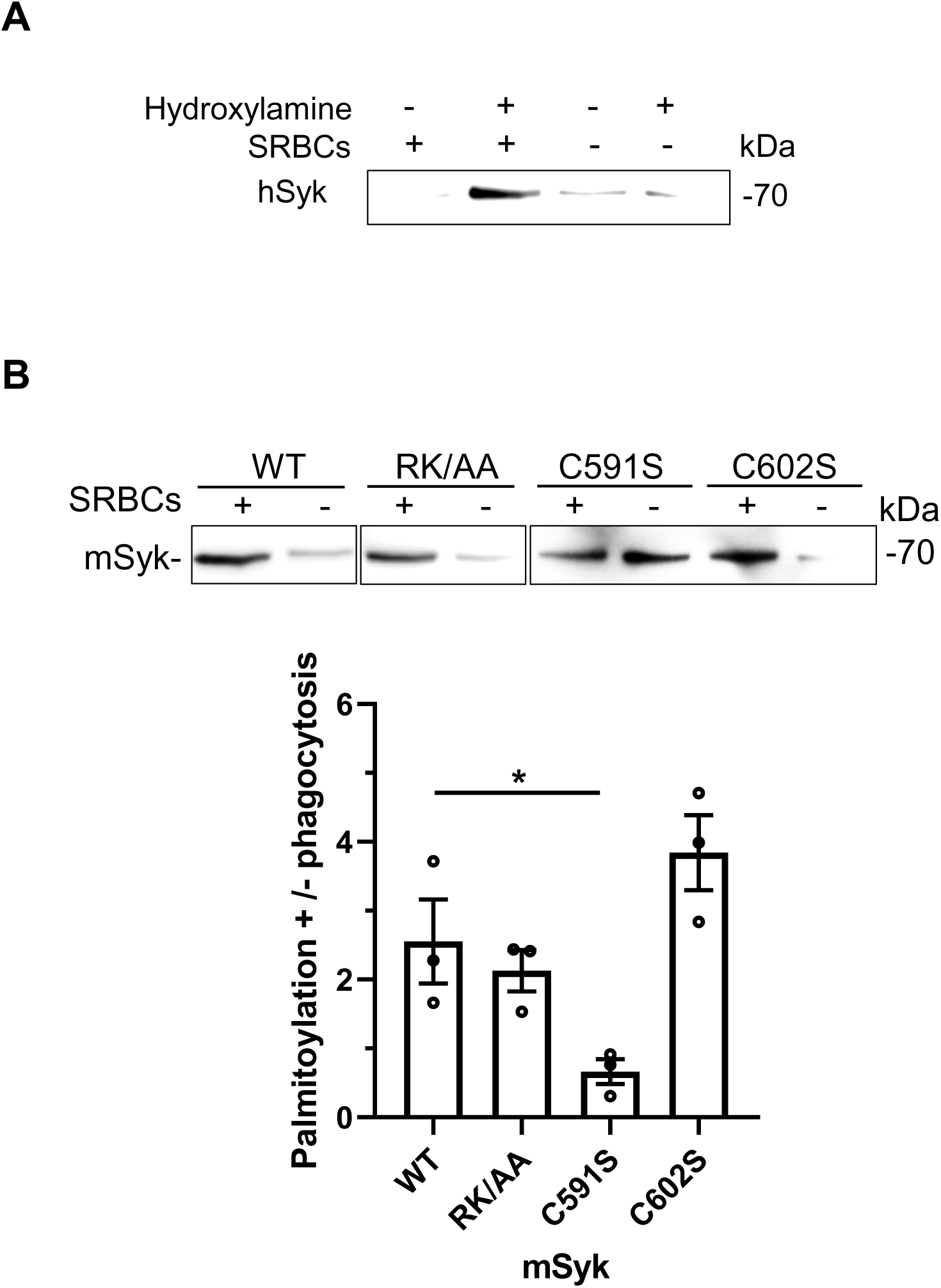
Syk is palmitoylated on mSyk-Cys501. **A**, hMDMs were allowed to phagocytose opsonized SRBCs as indicated and cells lysates were treated with hydroxylamine then HPDP-Biotin to replace acyl groups by biotin, before purifying biotin-labeled proteins on streptavidin-agarose and Syk Western blot. **B**, ΔSyk RAW cells were transfected with the indicated version of mSyk before SRBCs phagocytosis as indicated, acyl-biotin exchange using hydroxylamine and Syk Western blot. The graph shows the quantification from 3 independent experiments (mean ± SEM). One Way ANOVA (*, p<0.05).

### DDHC5 is responsible for Syk palmitoylation

The results showing that Syk is palmitoylated upon phagocytosis only (Figure 2) indicate that the enzyme responsible for its palmitoylation is present at the plasma membrane. When we examined the intracellular localization of the different DHHC enzymes in RAW macrophages, we observed that, in agreement with our previous study on PC12 neuroendocrine cells (Chopard *et al*., 2018), DHHC5 and DHHC20 only were present at the plasma membrane (Figure S3). It has been shown that DHHC20 palmitoylates and thereby regulates the activity of DHHC5 (Plain *et al*, 2020). We first used siRNAs to deplete these DHHC enzymes in RAW macrophages. The siRNAs against DHHC5 depleted ∼50 % of their DHHC5 in RAW cells but conversely increased by 80% their DHHC20 level (Figure 3 A-B). On the other hand, siRNAs against DHHC20 reduced its level by ∼60% without significantly affecting the DHHC5 level.

**Figure 3.**
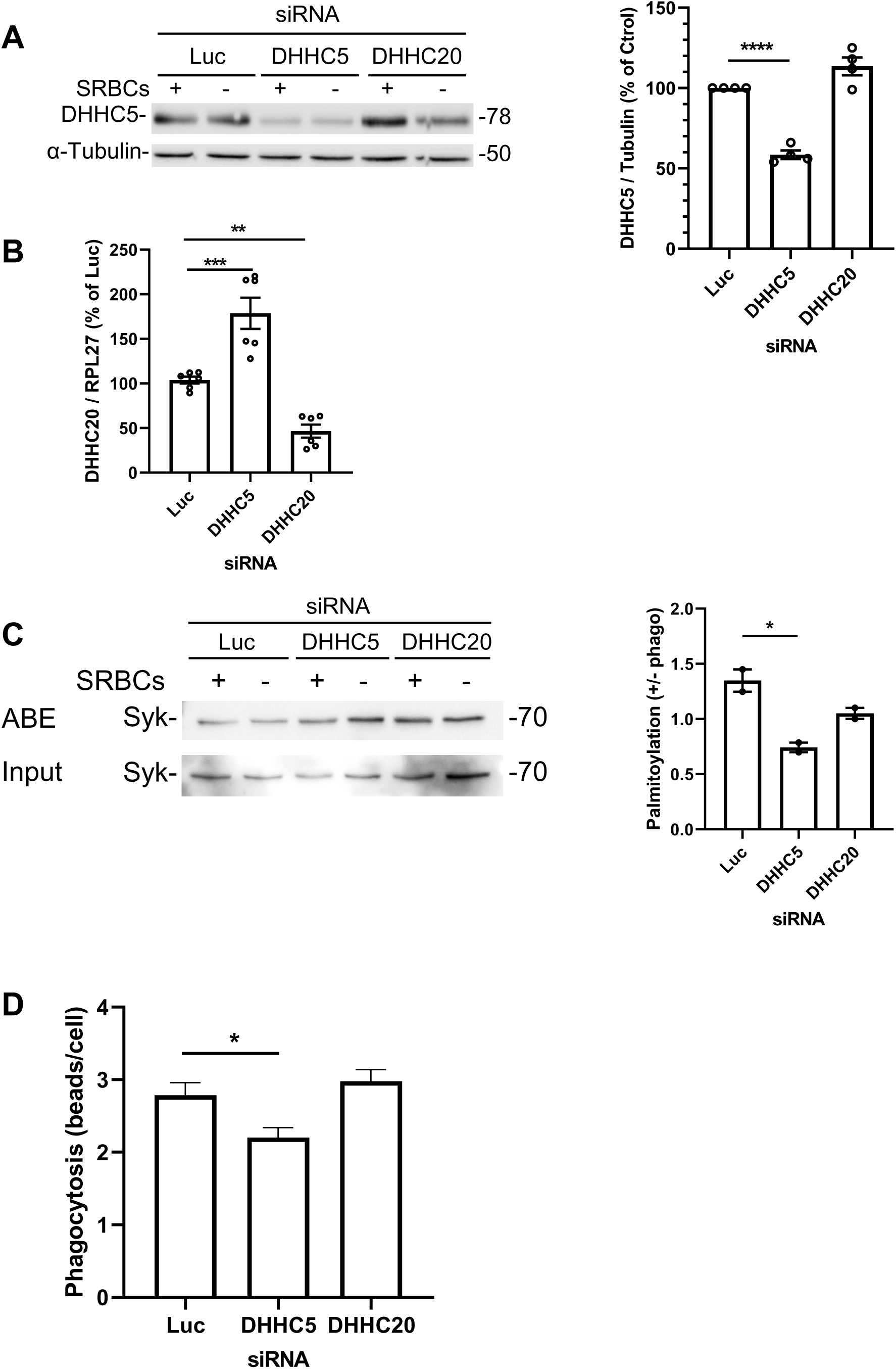
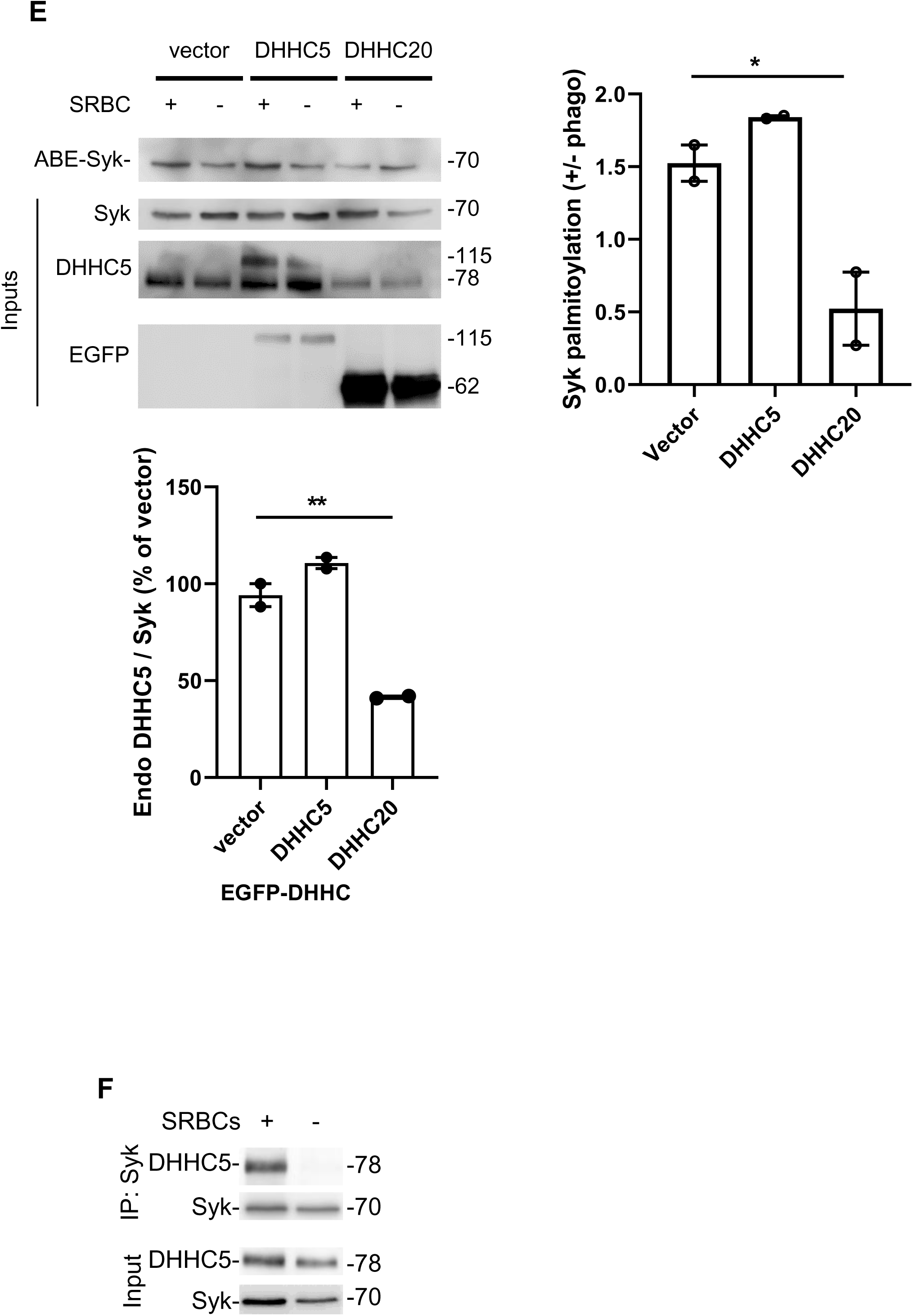
Syk is palmitoylated by DHHC5 and interacts with DHHC5. **A**, RAW cells were transfected with the indicated siRNA before phagocytosis of opsonized SRBCs for 5 min, cell lysis and Western blot against DHHC5 and tubulin. The graph shows the quantification from 4 experiments (mean ± SEM). **B**, RNA was extracted from transfected cells before qPCR for DHHC20 and RLP27. Means ± SEM (n=6). **C**, Syk palmitoylation is performed by DHHC5. Transfected RAW cells were allowed to phagocytose opsonized SRBCs before acyl-biotin exchange, purification on streptavidin-agarose and Syk Western blot. The graph shows the quantification (n=2 independent experiments). **D**, Depletion of cells in DHHC5 inhibits phagocytosis. RAW cells were transfected with the indicated siRNAs before assaying phagocytosis efficiency (mean ± SEM, n>178 transfected cells/condition). **E**, Effect of DHHC5 or DHHC20 overexpression on Syk palmitoylation. RAW cells were transfected with the indicated EGFP-mDHHC before phagocytosis of opsonized SRBCs, cell lysis and Syk palmitoylation assay using ABE. EGFP- mDHHC5 was detected at 115 kDa using anti-DHH5 and anti-EGFP, while endogenous mDHHC5 is at 78 kDa. EGFP-mDHHC20 (62 kDa) could only be detected using anti-EGFP. The graphs show the efficiency of Syk palmitoylation and the effect of DHHC overexpression on the endogenous DHHC5 level. One Way ANOVA (*, p<0.05; **, p<0.01; ***, p<0.001; ****, p<0.0001). **F,** DHHC5 interacts with Syk upon phagocytosis. hMDMs were allowed to phagocytose opsonized SRBCs, before Syk immunoprecipitation and Syk and DHHC5 western blots.

When we examined the effect of these siRNAs on Syk palmitoylation, we observed that siRNAs against DHHC5 significantly decreased Syk palmitoylation, while siRNAs against DHHC20 had no effect (Figure 3 C). Similar results were obtained when we quantified the effect of these siRNAs on phagocytosis efficiency (Figure 3D). These data are consistent with the identification of DHHC5 as the enzyme palmitoylating Syk, but do not exclude that DHHC5 could also palmitoylate other proteins essential for phagocytosis as well.

To confirm data from DHHC depletion using siRNAs, we used DHHC overexpression experiments. Expression of EGFP-mDHHC5, at level comparable to the endogenous mDHHC5, did not significantly increase the level of Syk palmitoylation, while overexpression of EGFP-mDHHC20 decreased Syk palmitoylation by 3-fold (Figure 3E). This is probably because EGFP-mDHHC20 overexpression decreased endogenous DHHC5 level by ∼60 %, a drop that could be due to overpalmitoylation of mDHHC5.

Interestingly, upon Syk immunoprecipitation from hMDMs, we observed using that DHHC5 specifically associates with Syk upon phagocytosis (Figure 3F). Altogether these results show that DHHC5 is responsible for Syk palmitoylation.

### mSyk is sulfenylated on Cys602

Syk is palmitoylated on its penultimate Cys (C591 for mSyk) and this modification is required for Syk to sustain phagocytosis (Figure 1B), but the role of the last Cys (Cys602 for mSyk) that is strictly needed for phagocytosis remained unclear. Syk contains in its C-terminal sequence a redox-sensitive motif previously identified in several non-receptor tyrosine kinases such as Src and Zap70. The Cys within this motif was found to be sulfenylated in the Syk-family kinase Zap70 (Thurm *et al*., 2017).

We first examined if Syk is sulfenylated using hMDMs and the specific reagent for Cys-SOH termed DCP- Bio1 (Hourihan *et al*, 2016) that enables to label sulfenylated Cys with biotin. Syk was purified by immunoprecipitation from DCP-Bio1- treated extracts, and biotin staining revealed a large number of bands, including the one corresponding to hSyk. Quantification confirmed that Syk was sulfenylated and that its sulfenylation level was decreased by 20% upon phagocytosis induction (Figure 4A). When we used ΔSyk RAW macrophages transfected with mSyk we also observed that Syk was desulfenylated upon phagocytosis. DCP-Bio1 labeling produced faint bands but quantification of 4 independent experiments showed that, unlike WT mSyk, neither Syk RK/AA, nor C591S or C602S were desulfenylated upon phagocytosis (Figure 4B). We thus concluded that Syk is desulfenylated upon phagocytosis and that PIP3 binding and palmitoylation are required for Syk desulfenylation. The sulfenylation site is most likely Cys602 since Cys591 is palmitoylated.

**Figure 4.**
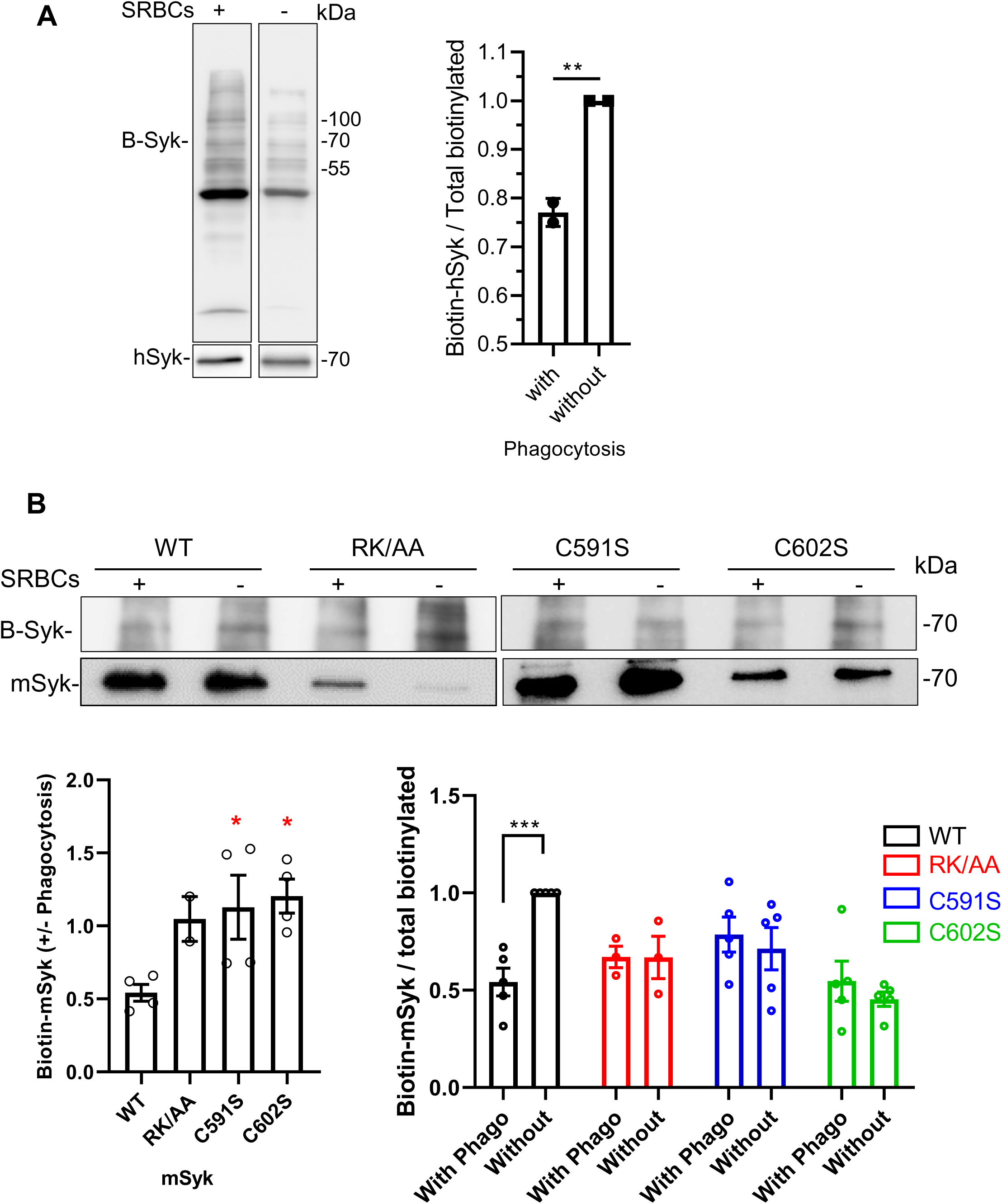
Syk is desulfenylated upon phagocytosis induction. **A**, hMDMs were allowed to phagocytose opsonized SRBCs as indicated and cells lysates were treated with DCP-Bio1 to attach biotin to sulfenylated Cys. After Syk immunoprecipitation and Western blot, biotin-hSyk (B-Syk) was detected using Extravidin- peroxidase. The graph shows the quantification of biotin-hSyk (mean ±SEM, n=2). Unpaired Student’t Test (**, p<0.01). **B**, mSyk is sulfenylated on Cys602. ΔSyk RAW cells were transfected with the indicated mSyk mutant before phagocytosis of opsonized SRBCs. Cells lysates were treated with DCP-Bio1 before mSyk immunoprecipitation, Western blot and biotin staining. The graph shows the quantifications of biotin-Syk (mean ±SEM, n=4). One-Way or Two-Way ANOVA (*, p<0.05; ***, p<0.001).

To examine the effect of sulfenylation on the Syk structure, we performed molecular dynamics simulations. We observed that the desulfenylation of Cys602 increases the mobility of a flexible loop (residues 267-325) within the interdomain-B region (Figure 5A). Moreover, the loop movements were more oriented toward the outside of the WT Syk protein (Figure 5B). These results are consistent with a previous study of the Syk activation process that identified the Tyr348/352 residues within the flexible loop as crucial tyrosine residues whose phosphorylation by the Lyn Src-family kinase initiates Syk activation (Mansueto *et al*., 2019). Hence Syk desulfenylation likely facilitates its interaction and phosphorylation by Src family kinases by enhancing the mobility and the exposure of this loop.

**Figure 5.**
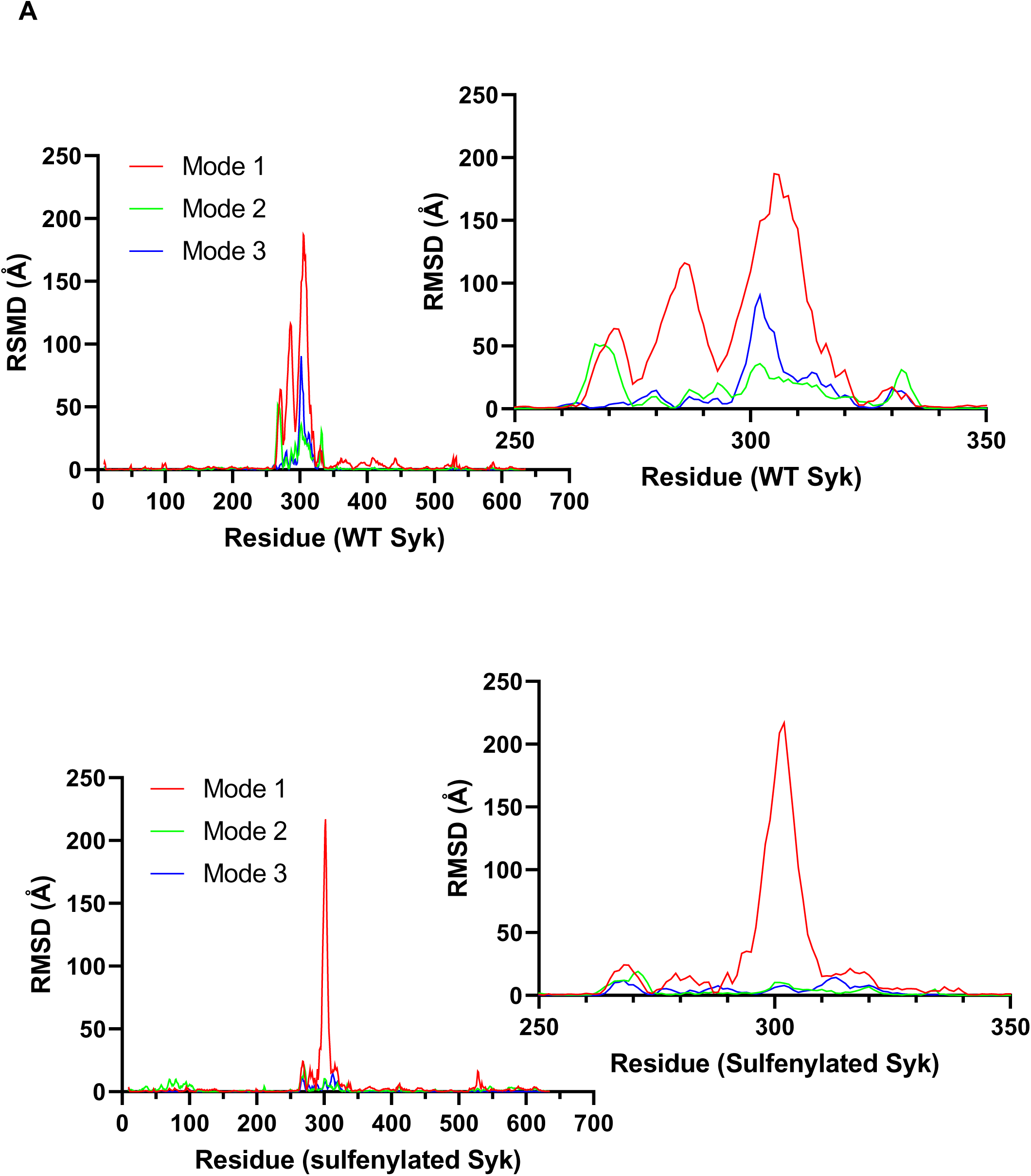
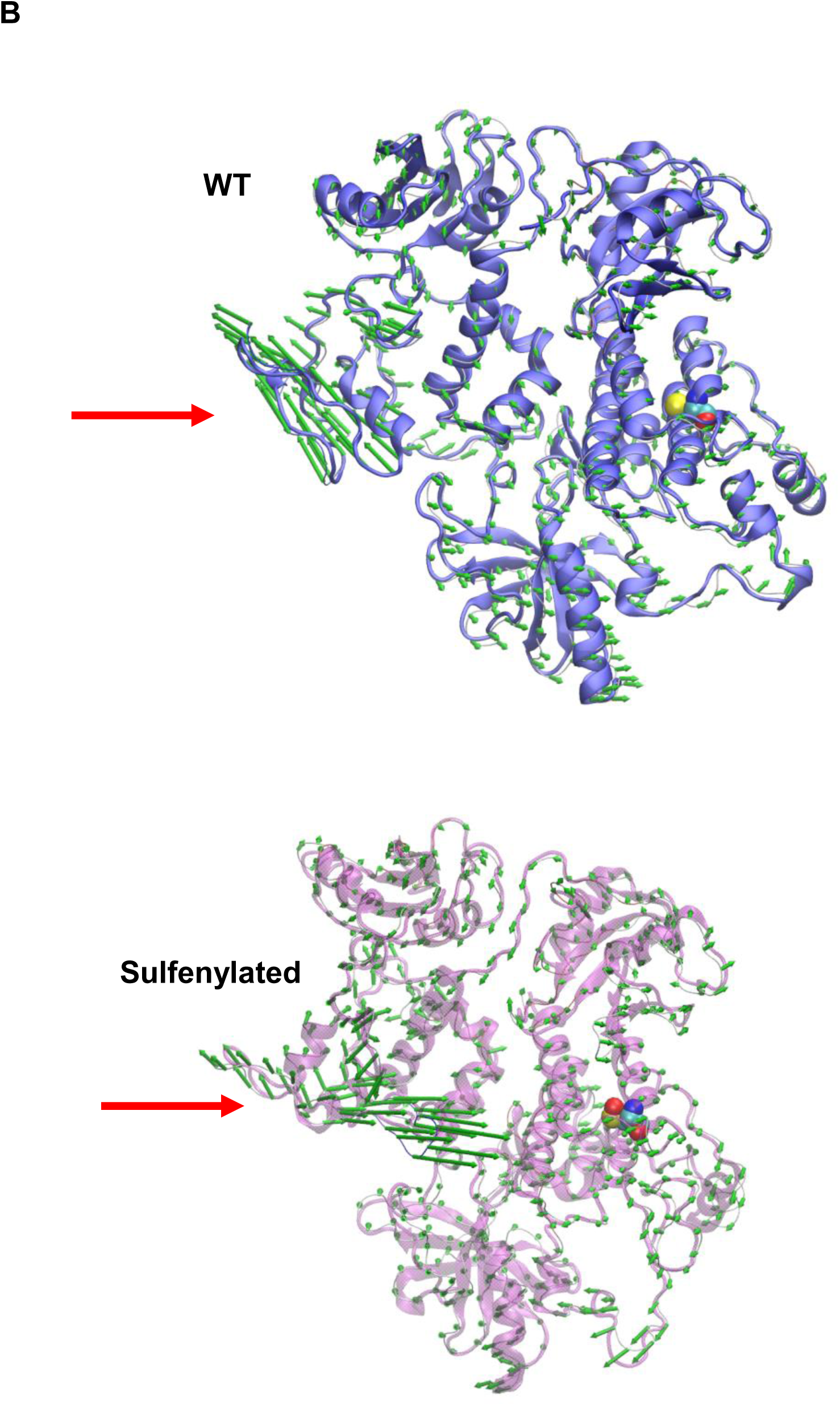
Molecular dynamics studies indicate that hSyk desulfenylation induces a higher mobility and exposure of a loop that includes the 263–335 residues. **A**, Mobility of the protein backbone (RMSF, Å) in 3 different modes. **B**, mobility of the protein. Arrows point to the flexible loop. For the WT the loop is moving toward the outside of the protein, while for sulfenylated hSyk, movements are smaller and directed toward the center of the molecule.

### Sulfenylation, palmitoylation and binding to PIP3 are required for Syk phosphorylation and activation

We then monitored the consequences of sulfenylation, palmitoylation and binding to PIP3 on Syk activation by following its autophosphorylation on two key Tyr residues (525/526 for hSyk) within Syk activation loop, in the kinase domain. Their phosphorylation was found to be a prerequisite for Syk activation (Zhang *et al*, 2000). Using the corresponding Syk mutants we found that catalytic activity, SH2 binding, desulfenylation, palmitoylation and PIP3 binding are all required for Syk activation (Figure 6).

**Figure 6.**
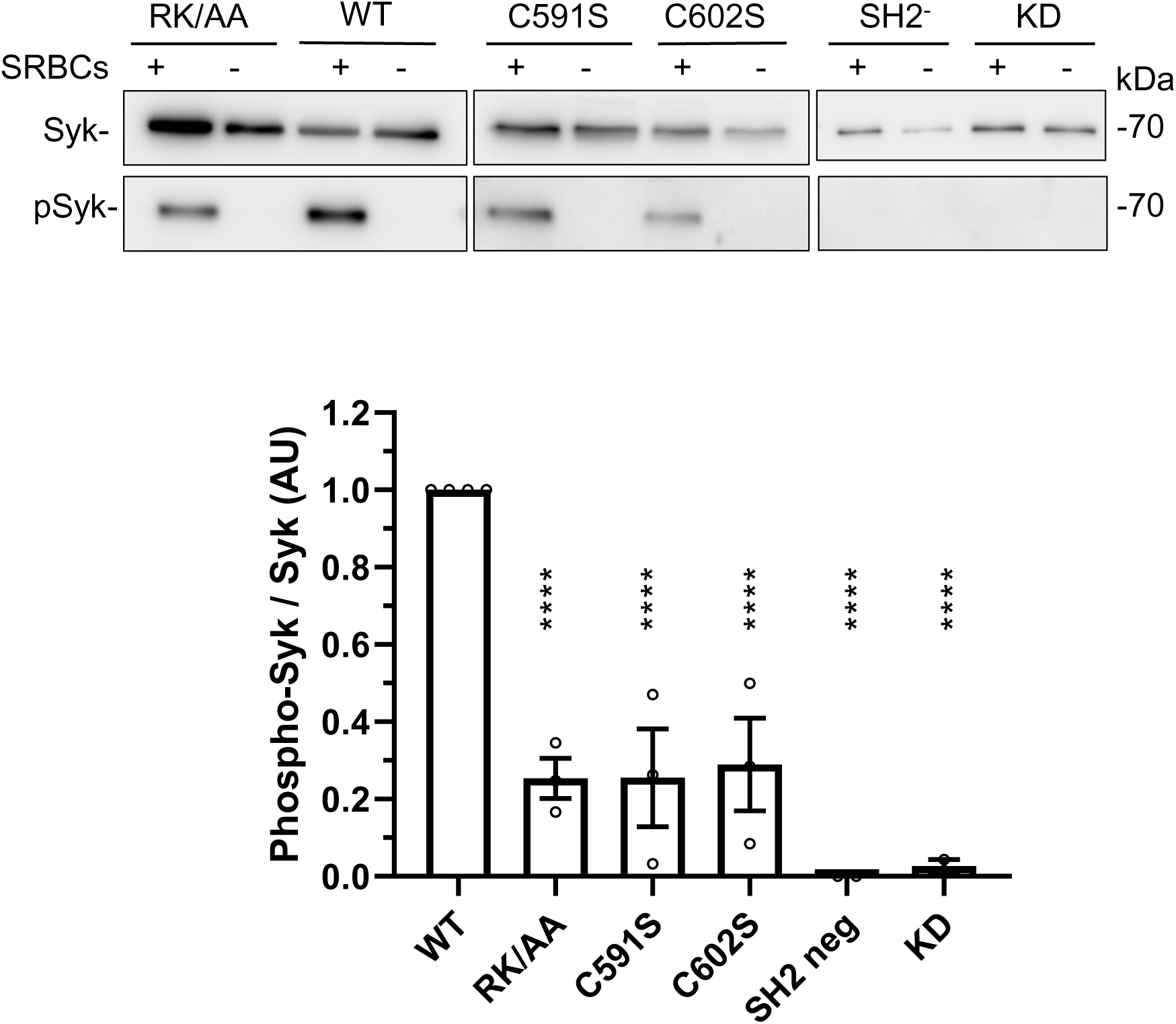
Sulfenylation, palmitoylation and PIP3-binding of Syk are required for efficient Syk phosphorylation upon phagocytosis. ΔSyk RAW cells were transfected with the indicated mSyk mutant before phagocytosis of opsonized SRBCs, SDS-PAGE, and Western blot using an anti-phosphoTyr-mSyk 519/520 (equivalent to Tyr 525/526 of hSyk). The graph shows the quantification of n=4 independent experiments (mean ± SEM). One Way ANOVA; ****, p<0.0001.

### Syk sulfenylation / desulfenylation are required for Syk catalytic activity

We then examined the effects of sulfenylation, palmitoylation and binding to PIP3 on Syk catalytic activity. To this end we used an *in cellulo* FRET assay (Xiang *et al*, 2011). We first monitored the kinetics of Syk activation following the onset of phagocytosis. In agreement with the previously reported kinetics of Syk phosphorylation (Raeder *et al*., 1999), we observed that Syk catalytic activity is maximal after 5 min of phagocytosis (Figure 7A). When we assessed the catalytic activity of the different Syk mutants we found that the C602S was inactive while Syk-591S and Syk-RK/AA were catalytically active (Figure 7B). These results thus indicated that sulfenylation/ desulfenylation is required for Syk catalytic activity while PIP3 binding and palmitoylation are not necessary. Control mutants showed that Syk catalytic site and SH2-binding are both critical for Syk kinase activity *in cellulo*.

**Figure 7.**
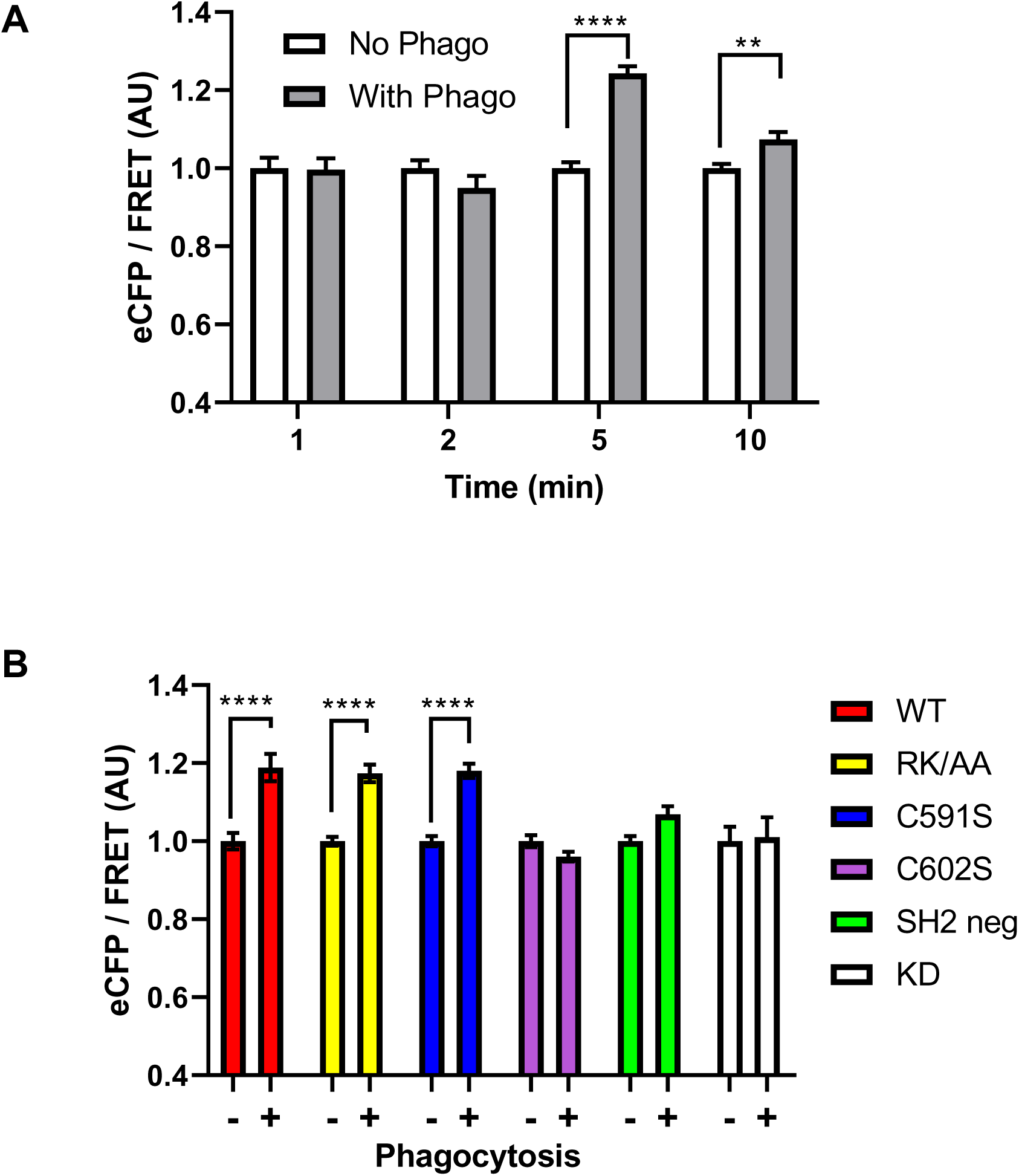
Syk sulfenylation but not palmitoylation or PIP3 binding is required for Syk catalytic activity. **A**, RAW cells were transfected with the FRET Syk biosensor before phagocytosis of opsonized SRBCs for the indicated time. After fixation, cells (n>50) were imaged using a confocal microscope to monitor ECFP and FRET fluorescence. Results were normalized using values for 1 min time. **B**, ΔSyk RAW cells were transfected with the FRET Syk biosensor and the indicated mSyk mutant before phagocytosis of opsonized SRBCs for 5 min. After fixation, cells (n>30) were imaged using a confocal microscope. Two Way ANOVA; **, p<0.01; ****, p<0.0001.

### Syk palmitoylation is important for its localization at the phagocytic cup

Because palmitoylation usually stabilizes the association of proteins with membranes (Ko & Dixon, 2018), we assessed whether Syk palmitoylation affected its localization at the phagocytic cup. To this end, we used RAW macrophages transfected with EGFP-Syk. As previously observed in B cells (Ma *et al*, 2001), Syk is essentially cytosolic and recruited to the cell membrane upon stimulation. We followed Syk accumulation at the phagocytic cup using line plots across the cell and compared Syk accumulation at the cup with that on the opposite region of the plasma membrane (Figures 8 and S4). Statistical analysis showed that Syk was ∼4-fold more concentrated at the cup compared to control areas of the plasma membrane (Figure 8). Non-palmitoylable Syk (C591S) was only 1.9-fold concentrated at the cup, while non-sulfenylable Syk (C602S) and Syk-RK/AA unable to bind PIP3 behaved as WT Syk. Control Syk mutants with inactivated SH2 domains or catalytic site were not concentrated at the cup. These results indicated that Syk catalytic activity, P-ITAM binding and palmitoylation are all needed to insure Syk localization at the phagocytic cup, but sulfenylation/ desulfenylation is not required.

**Figure 8.**
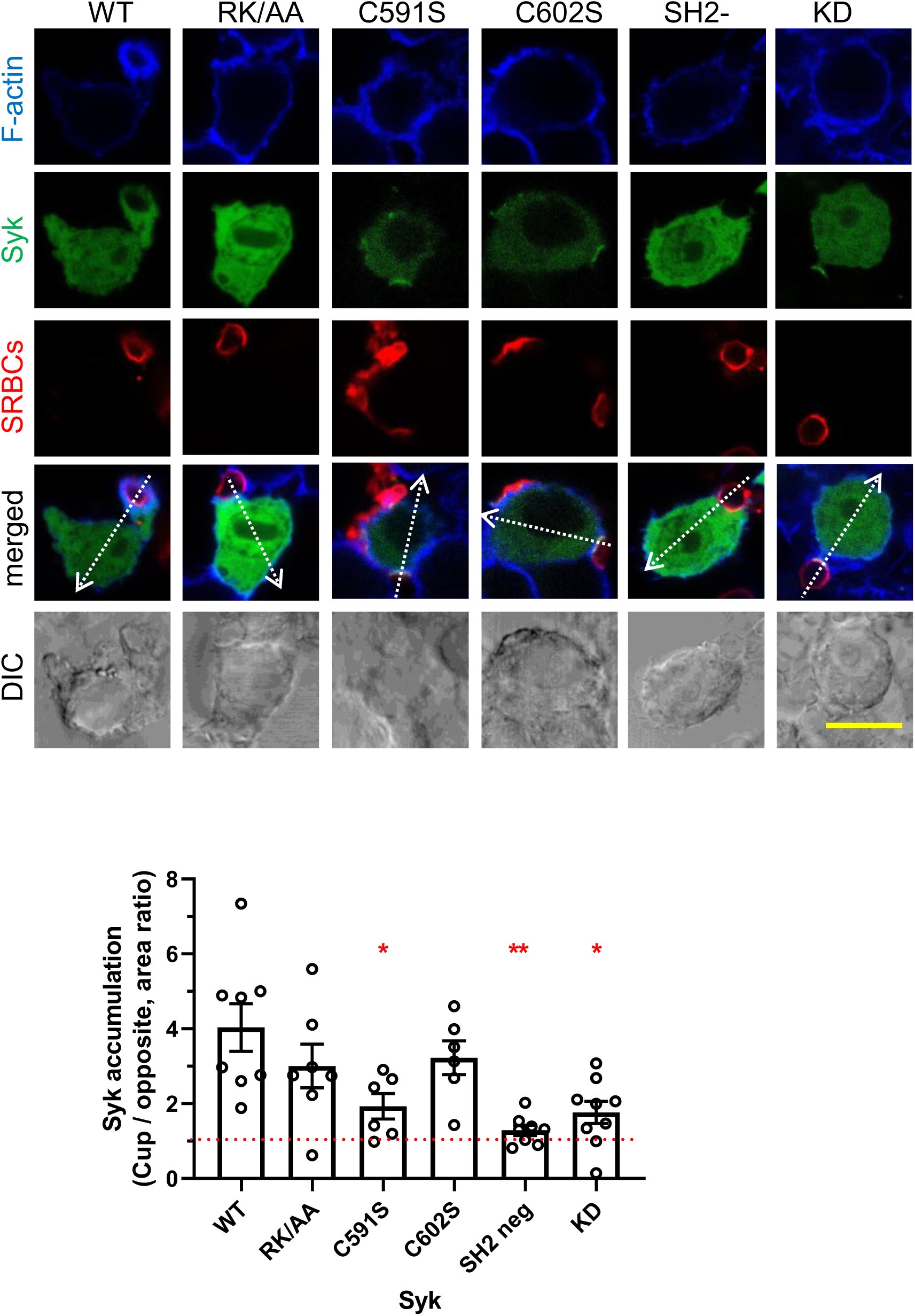
Syk palmitoylation is required for its recruitment at the phagocytic cup. ΔSyk RAW cells were transfected with the indicated EGFP-mSyk mutant before phagocytosis of opsonized SRBCs for 5 min. Cells were then fixed and stained with phalloidin. Line plots (width: 10 pixels) showed on the merged images enabled to follow signal accumulation at the phagocytic cup. Bar, 10 µm. The graph shows the ratio (mean ± SEM) of the fluorescence peaks cup / opposite side of the cell, from n=6-9 cells. One Way ANOVA; *, p<0.05; **, p<0.01.

### Syk palmitoylation regulates the recruitment of Cdc42 but not Rac1 to the phagocytic cup

The Rho-family GTPases Rac1 and Cdc42 are key regulators of the actin polymerization process that enable pseudopod extension during phagocytosis (Freeman & Grinstein, 2014; Mylvaganam *et al*., 2021). We examined the role of Syk modifications on the recruitment at the cup of these two GTPases. WT Syk enabled Rac1 and Cdc42 recruitment at the cup, with a 3-fold average enrichment (Figures 9A-B, S5 and S6). No GTPase recruitment was observed for C602S, SH2-deficient and kinase-dead Syk mutants, indicating that sulfenylation/desulfenylation, catalytic activity and P-ITAM binding were required for Syk to trigger Rac1 and Cdc42 recruitment at the cup. While the Syk RK/AA mutant unable to bind PIP3 behaved as the WT, the only mutant that differentially affected the recruitment of the GTPases was the palmitoylation deficient mutant C591S that enabled the recruitment of Rac1 (Figures 9A and S5) but not Cdc42 (Figures 9B and S6).

**Figure 9.**
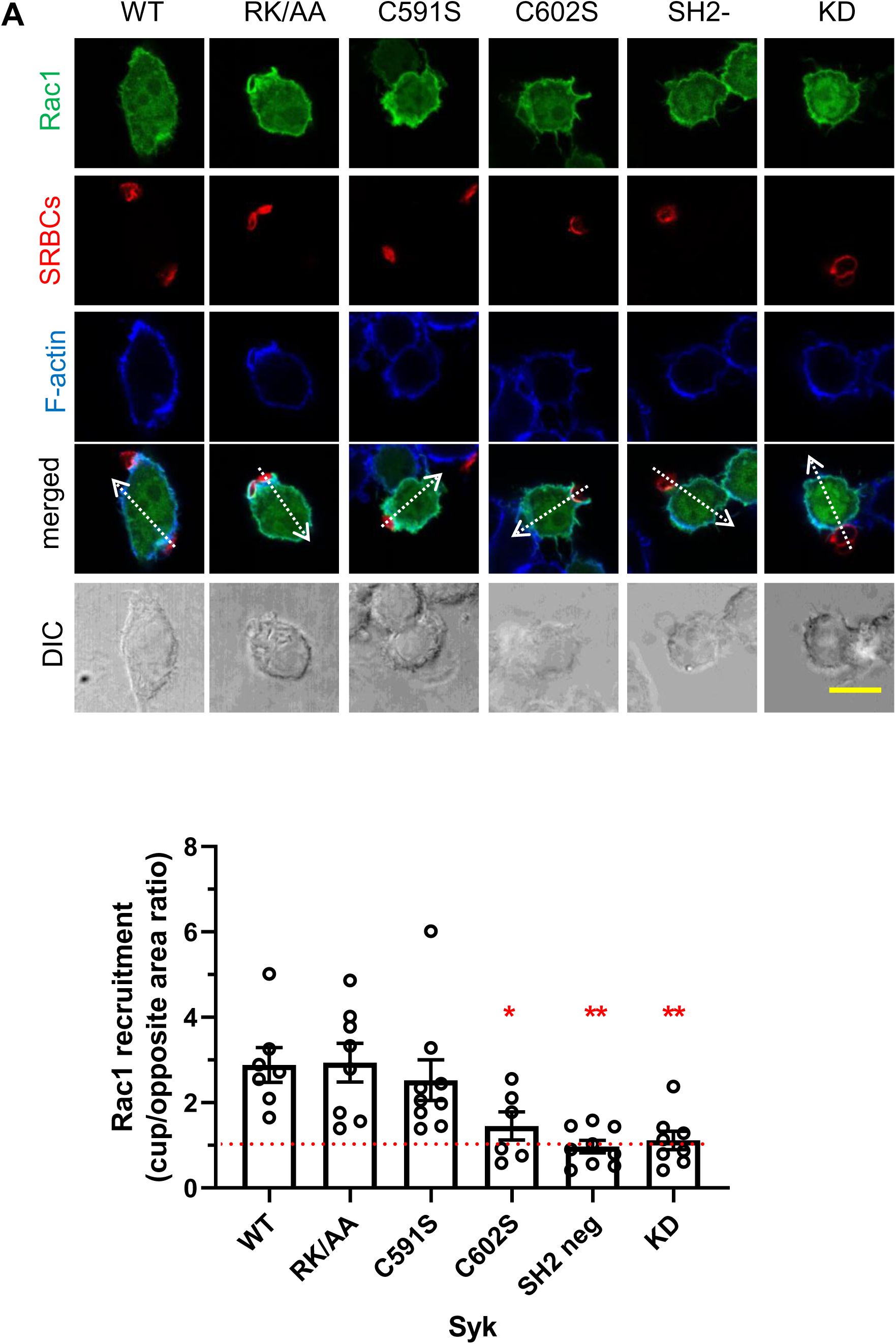
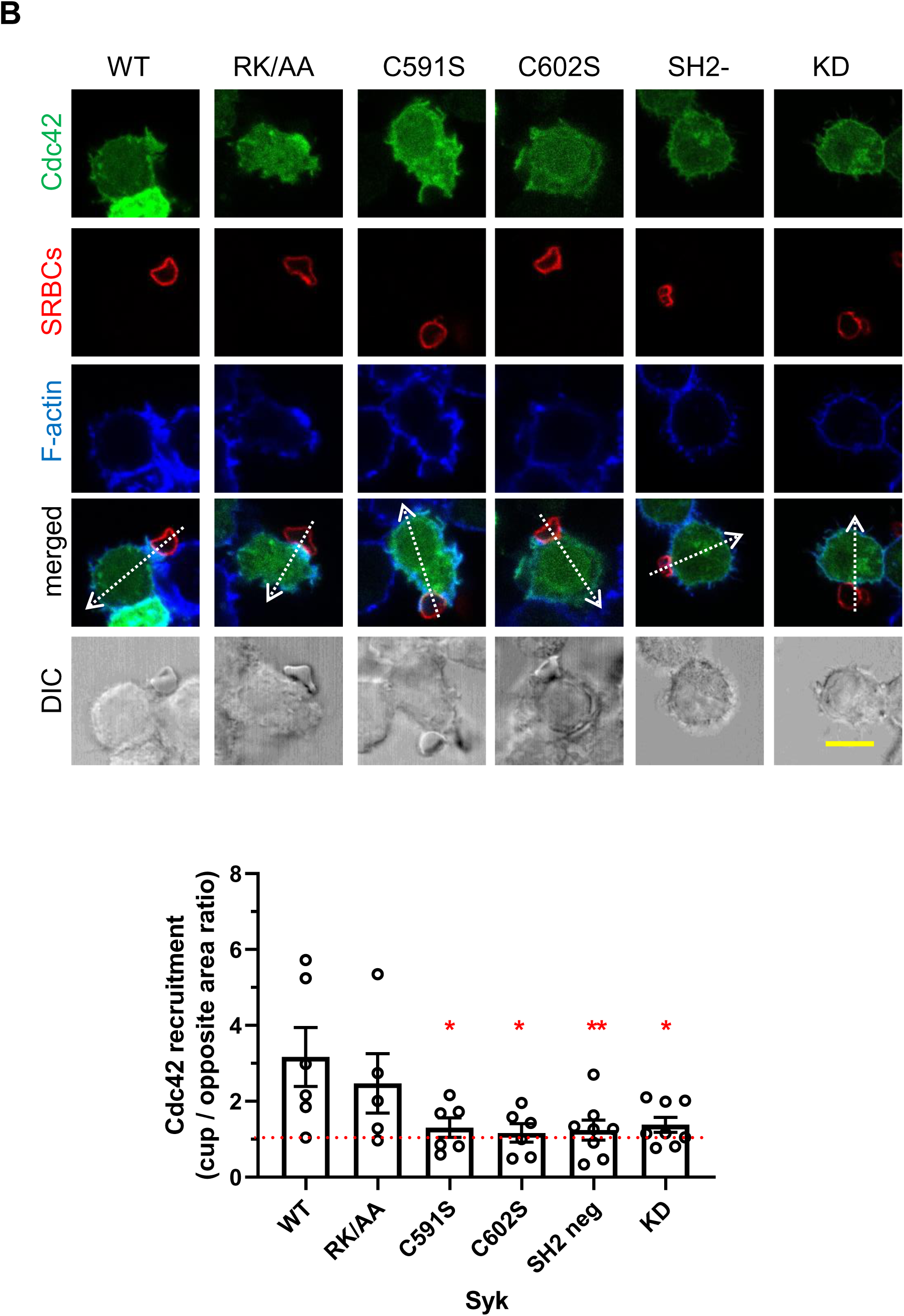
Syk palmitoylation is required for the recruitment of Cdc42 but not of Rac1 at the phagocytic cup. ΔSyk RAW cells were transfected with vectors encoding for (**A**) EGFP-Rac1 or (**B**) EGFP-Cdc42, and the indicated FLAG-mSyk mutant before phagocytosis of opsonized SRBCs for 5 min. Cells were then fixed and stained with phalloidin. Line plots (width: 10 pixels) showed on the merged images enabled to follow signal accumulation in the phagocytic cup. Bars, 10 µm. The graphs show the ratio (mean ± SEM) of the areas of the fluorescence peaks at the cup / opposite side of the cell, from n=6-9 cells. One Way ANOVA; *, p<0.05; **, p<0.01.

### Syk sulfenylation/desulfenylation and palmitoylation are required for PIP3 and DAG production at the phagocytic cup

Syk is required for PIP3 and DAG production at the phagocytic cup (Freeman & Grinstein, 2014; Mylvaganam *et al*., 2021). We followed the production of these second messengers using fluorescent chimeras. The different Syk mutants similarly affected the production of both PIP3 and DAG (Figures 10A and S7). While sulfenylation/desulfenylation, SH2-deficient and kinase-dead mutants did not allow significant production of these second messengers, the non-palmitoylable Syk mutant was ∼50 % less efficient than WT Syk in inducing PIP3 and DAG production. Results with the RK/AA mutant indicated that Syk binding to PIP3 is not required for the production of these messengers.

**Figure 10.**
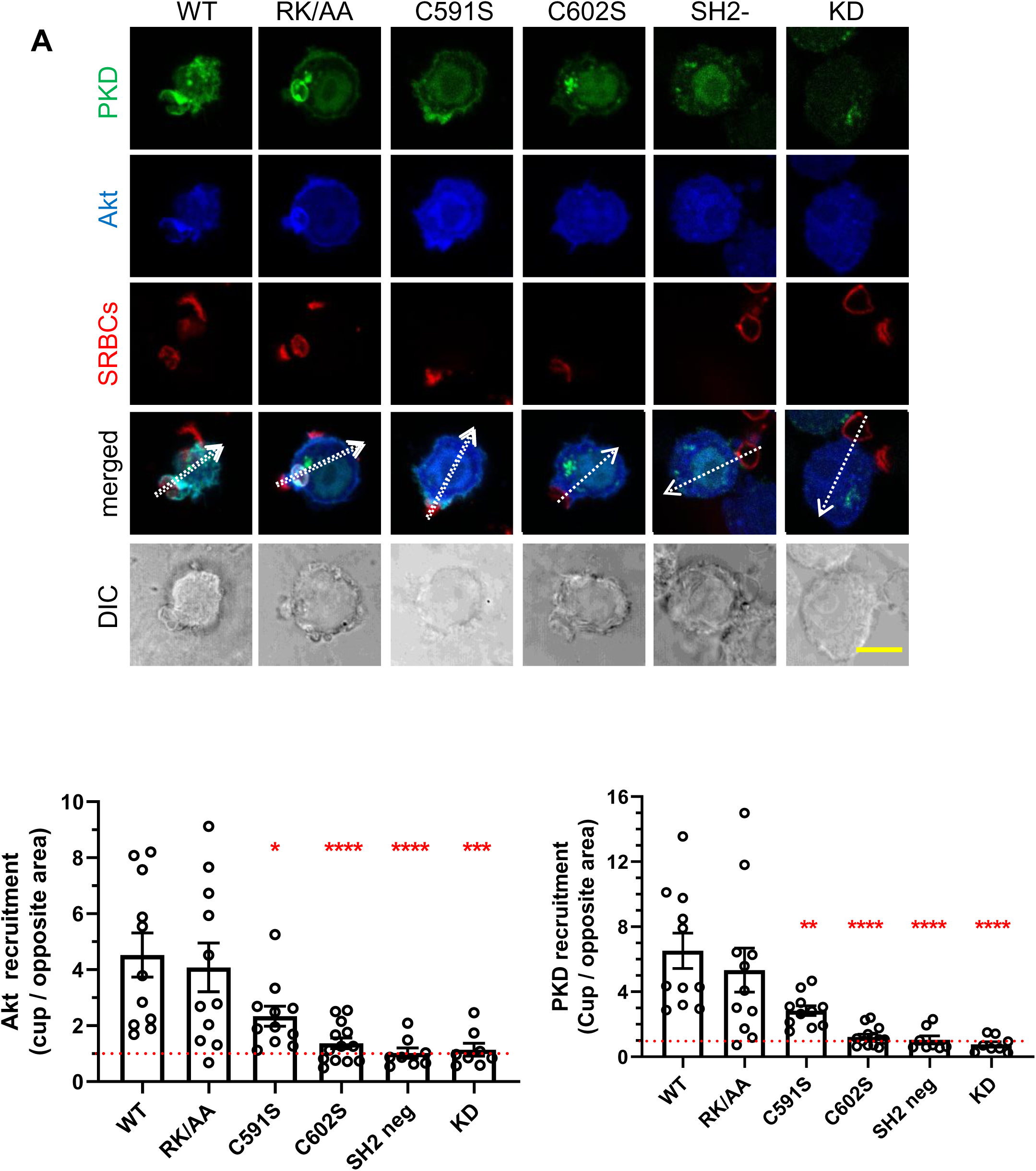
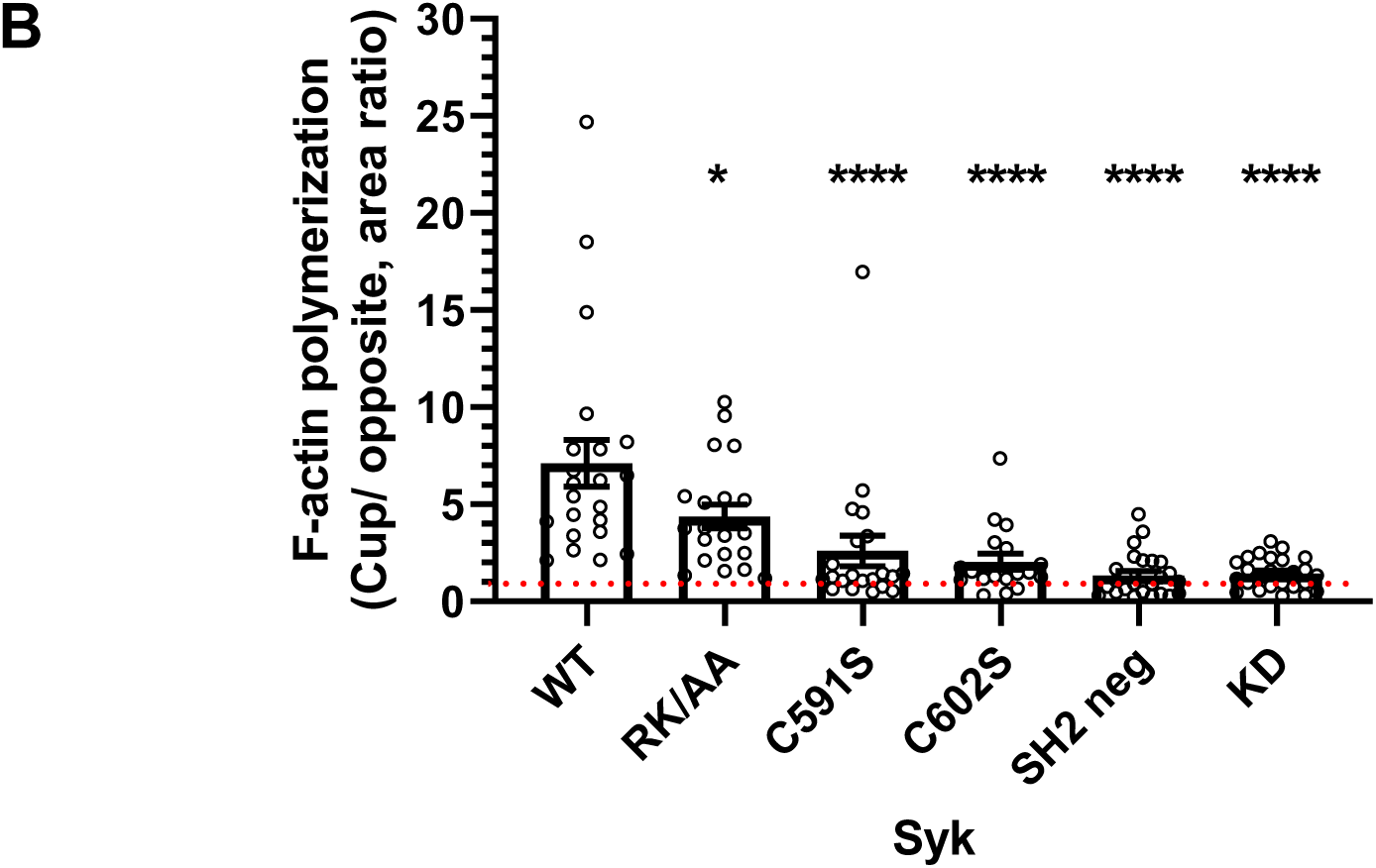
Syk sulfenylation and palmitoylation are required for PIP3 and DAG production, and for F-actin polymerization at the phagocytic cup. **A**, ΔSyk RAW cells were transfected with vectors encoding for EGFP- PKD-C1ab, mCherry-Akt and the indicated FLAG-mSyk mutant as indicated before phagocytosis of opsonized SRBCs for 5 min. Cells were then fixed and stained with phalloidin. Line plots (width: 10 pixels) showed on the merged images enabled to follow signal accumulation in the phagocytic cup. Bar, 10 µm. The graph shows the ratio (mean ± SEM) of the fluorescence peaks cup / opposite side of the cell, from n=8-12 cells. **B**, F-actin accumulation at the cup was measured for the indicated mutants using fluorescent phalloidin and 18-25 cells. One Way ANOVA; *, p<0.05; **, p<0.01;***, p<0.001; ****, p<0.0001.

### All Syk modifications are required to allow actin polymerization at the phagocytic cup

F-actin polymerization at the phagocytic cup is driving pseudopod extension and phagocytosis (Freeman & Grinstein, 2014; Mylvaganam *et al*., 2021). We found that all Syk mutants were affected in their capacity to initiate F-actin polymerization at the cup (Figure 10B). Not surprisingly, their capacity to initiate F-actin accumulation at the cup (Figure 10B) essentially corresponded to their ability to sustain phagocytosis (Figure 1B). While the PIP3-binding deficient mutant showed an intermediate phenotype, the mutants deficient in sulfenylation/desulfenylation, P-ITAM binding or catalytic prevented almost entirely F-actin concentration at the cup and phagocytosis. The non-palmitoylable mutant, despite the fact that it allowed only minute amounts of actin accumulation at the cup (Figure 10B), enabled significant phagocytosis (∼30% of WT, Figure 1B).

### An updated model for Syk activation during phagocytosis

Based on our results (summarized in Table 1) we propose the following model for the activation of Syk during phagocytosis (Figure 11). The first step consists of the Syk recruitment at the phagocytic cup by P- ITAMs. This initial membrane-targeting allows Syk to encounter and interact with DHHC5, which is a membrane protein (Chopard *et al*., 2018). The non-palmitoylable mutant does not stay at the cup (Figure 8) but is nevertheless able to initiate Rac1 recruitment (Figure 9A). Syk palmitoylation secures its association to the cup and allows more PIP3 production and consequently more significant PIP3 binding. Palmitoylation is also needed for Cdc42 recruitment at the cup.

**Table I.** Summary of the mSyk-mutant phenotypes. The C591S mutant is non-palmitoylable, the C602S non sulfenylable and the RK/AA mutant (R219A/K221A) unable to bind PIP3. Residue numbering refers to mSyk.

| **mSyk Mutant** | **C591S** | **C602S** | **RK/AA** |
| --- | --- | --- | --- |
| Palmitoylation | - | + | + |
| Sulfenylation / desulfenylation | - | - | +/- |
| mSyk phosphorylation on Tyr 519/520 | - | - | - |
| Catalytic activity *in cellulo* | + | - | + |
| mSyk recruitment at the phagocytic cup | - | + | + |
| Rac1 recruitment at the phagocytic cup | + | - | + |
| Cdc42 recruitment at the phagocytic cup | - | - | + |
| PIP3 and DAG production at the cup | - | - | + |
| Actin polymerization at the phagocytic cup | - | - | +/- |
| FcγR-mediated Phagocytosis | +/- | - | +/- |

**Figure 11.**
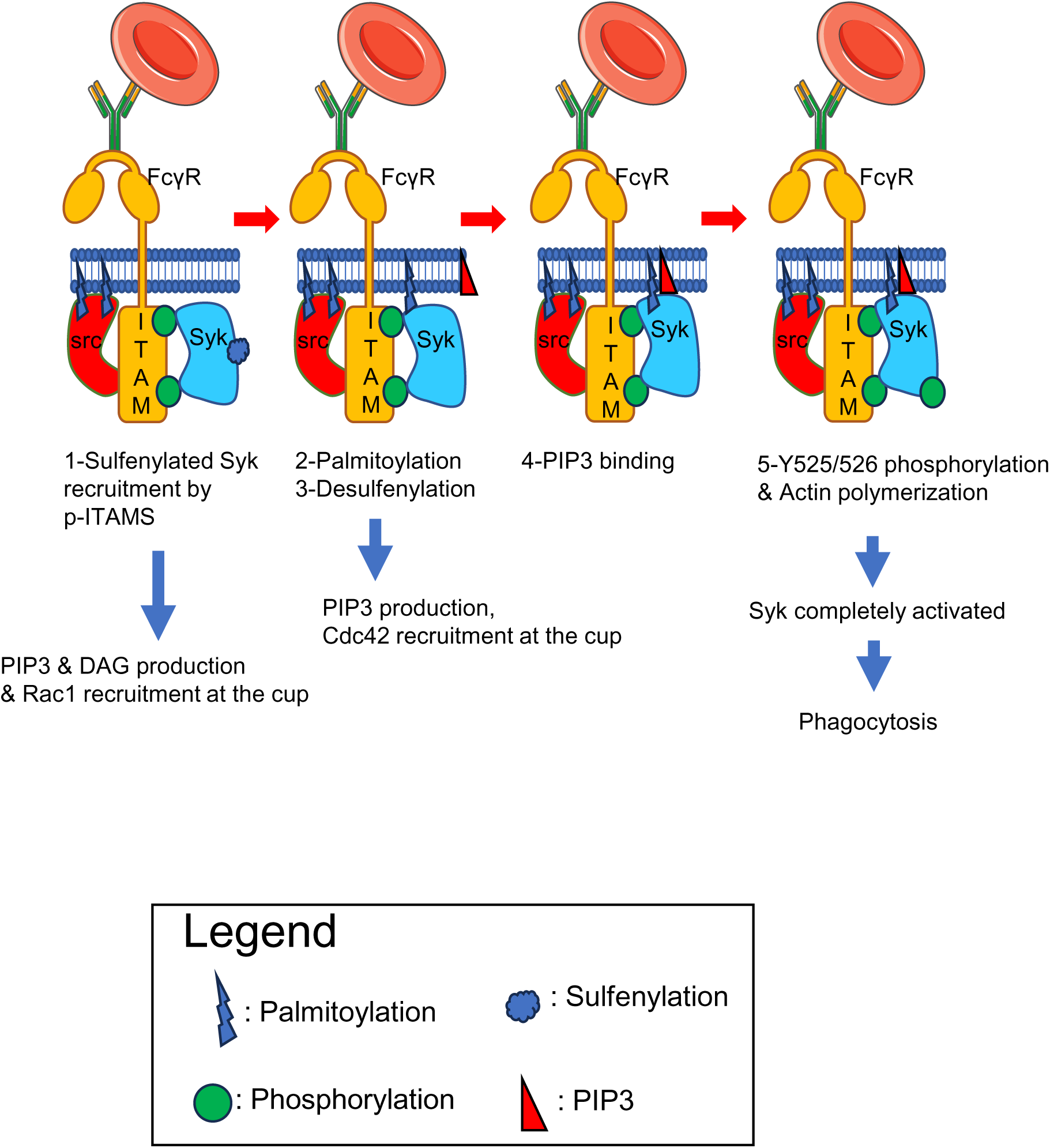
An updated model for Syk activation. Based on our results (Table I) we propose the following model for Syk activation. It involves 5 successive steps, 1-recruitment of Syk by P-ITAMs at the plasma membrane, 2-Syk palmitoylation, 3-Syk desulfenylation, 4-Syk-PIP3 binding and 5-Syk phosphorylation and actin polymerization. Rac1 takes place as soon as Syk is recruited by P-ITAMs at the plasma membrane, while Cdc42 recruitment requires Syk to be palmitoylated and desulfenylated.

Results obtained using the non-palmitoylable Syk mutant indicates that desulfenylation takes place after palmitoylation and Syk recruitment at the cup and that this is required for Syk catalytic activity. Cdc42 recruitment at the cup requires both palmitoylation and desulfenylation. The Syk mutant that is unable to bind PIP3 is not affected in the early steps of mSyk activation (Syk, Rac 1 and Cdc42 recruitment at the cup, PIP3 and DAG production) but it nevertheless failed to be phosphorylated on the key 519/520 Tyr residues involved in Syk activation (Figure 6) and to initiate F-actin polymerization (Figure 10B), indicating a crucial role of PIP3 binding in the late stages of Syk activation.

## Discussion

We here reveal two previously unknown post translational modifications of Syk, palmitoylation and sulfenylation / desulfenylation. Palmitoylation and desulfenylation were found to specifically take place during Syk activation upon FcγR-mediated phagocytosis in macrophages. A proteomic study of S-acylated proteins of RAW macrophages (Merrick *et al*, 2011) did not identify Syk because the macrophages were not stimulated. We identified mSyk-C591 (equivalent to hSyk-C597) as the palmitoylated residue. This residue is equivalent to Zap70-C564 that was suggested to be palmitoylated upon TCR stimulation (Tewari *et al*., 2021), a result that could not be confirmed in a later study (Schultz *et al*., 2022). Using both siRNA-based and overexpression approaches we observed that Syk palmitoylation is performed by DHHC5 that could be co-immunoprecipitated with Syk upon phagocytosis only. We showed that Syk palmitoylation by DHHC5 is regulated by DHHC20 that regulates DHHC5 intracellular level (this study) and palmitoylation (Plain *et al*., 2020). Palmitoylation is known to stabilize the association of cytosolic proteins once targeted to the membrane so that they can meet the appropriate DHHC enzyme (Chopard *et al*., 2018). Syk recruitment at the cup requires its binding to P-ITAMs (Figure 8) that enables its palmitoylation so that Syk remains at the cup and efficiently initiates downstream signaling such as Cdc42 recruitment, PIP3 and DAG production and, eventually, F-actin polymerization. Even in actively phagocyting macrophages Syk is largely a cytosolic protein (Figure 8) and only a minor fraction of Syk is recruited at the cup and palmitoylated.

Sulfenylation is another, albeit less studied, post-translational modification of Cys residues (Wages *et al*, 2015). We identified mSyk-Cys602 as a target of sulfenylation. Unlike palmitoylation, we observed that Syk is desulfenylated upon phagocytosis. Since the non-palmitoylable mutant (mSyk-C591S) is not desulfenylated, Syk desulfenylation likely takes place after palmitoylation, when Syk is already stably associated with the phagocytic cup. Molecular dynamics studies indicated that Syk desulfenylation enhances the mobility and the exposure of a loop formed by residues 267-325. Together with the binding of P-ITAM (Mansueto *et al*., 2019), desulfenylation that enhances the mobility and exposition of this loop within the interdomain-B of Syk likely favors the recognition and phosphorylation of nearby Syk-Tyr residues 348/352 by Src kinases.

Syk sulfenylation takes place on mSyk-Cys602 (hSyk-Cys608) within a redox motif that was observed to regulate Zap70 activity (Thurm *et al*., 2017). For both Syk and Zap70, mutation of the sulfenylated Cys resulted in decreased expression and catalytic activity. Since mutation of the sulfenylated Syk-Cys does not prevent Syk palmitoylation (Figure 2), the redox state of Syk does not affect palmitoylation, unlike CD81 for instance (Clark *et al*, 2004). Syk redox sensor is most likely responsible for the activation of Syk by H2O2 that was observed in B-cells (Schieven *et al*, 1993). Since the conformation and activity of Src kinases was also observed to be regulated by sulfenylation (Heppner *et al*, 2018), sulfenylation/ desulfenylation is likely a key regulator of phagocytosis efficiency.

Membrane association of Syk at the phagocytic cup and, most presumably, on ITAM-bearing receptors in general, is thus a complex process involving at least three different interactions. The initial recruitment on P- ITAMs is performed by Syk-SH2 domains, allowing palmitoylation. Since the mutant unable to bind PIP3 is palmitoylated, PIP3 binding likely takes place after palmitoylation. Syk recruitment at the cup enables PIP3 production, indicating that an amplification loop may exist, favoring Syk-PIP3 binding and further PIP3 production.

Rac1 and Cdc42 regulate F-actin polymerization at the phagocytic cup, but their role in pseudopod extension is different (Hoppe & Swanson, 2004; Massol *et al*, 1998). The molecular basis for this differential activation pattern is not clear, although it is probably linked to their recruitment by different phosphoinositides, i.e. PIP3 for Rac1 and PI(4,5)P2 for Cdc42 (Heo *et al*, 2006). We observed here that Syk differentially regulates the recruitment of these GTPases. Results on the non-palmitoylable Syk indeed showed that Rac1 recruitment at the phagocytic cup does not require Syk presence at this level, while Cdc42 recruitment requires the presence of Syk at the cup. These results indicate that Syk is involved in the differential recruitment and activation of these GTPases at the phagocytic cup.

In conclusion, we described here two novel post translational modifications of Syk that occur on its two last Cys residues. The penultimate Cys is palmitoylated and the last Cys is sulfenylated/desulfenylated. These modifications strongly affected Syk activation and activity during phagocytosis.

## Materials and methods

### Antibodies, plasmids oligos and chemicals

Please see the reagent list (Table S1).

### Plasmids

mSyk expression vector was obtained from Addgene (#50045). FLAG-mSyk was prepared by inserting a FLAG epitope (DYKDDDDK) upstream Syk coding sequence in pCi vector (Promega). To prepare EGFP- mSyk, mSyk was inserted downstream EGFP in pEGFP-C1 (Clontech). Mutations were introduced using Quikchange lightning site-directed mutagenesis kit (Agilent # 210518), and coding sequences were entirely sequenced to check mutations.

### Macrophages

RAW 264.7 mouse macrophages (termed RAW cells) were obtained and cultured according to the recommendations of the American Tissue Culture Collection. Cells were checked monthly for the absence of mycobacterial contamination. They were transfected with plasmids using Jet-Optimus (Ozyme, France) as recommended by the manufacturer, and harvested 24 h after transfection. For siRNA transfection, the RNAiMAX reagent (Thermo Scientific) was used as recommended by the manufacturer. A mix of 4 different siRNAs were used against DHHC5 and DHHC20. Cells were harvested 24h after transfection.

For the preparation of human monocytes, freshly drawn human blood was obtained from the local blood bank (Etablissement Français du Sang, Montpellier, agreement # 21PLER2019-0106). Peripheral blood mononuclear cells were prepared by density gradient separation on Ficoll–Hypaque (Eurobio) before isolating monocytes using CD14 microbeads (Miltenyi Biotec). Monocytes were then matured to macrophages by cultivation for 6–8 days in RPMI supplemented with 10 % FCS and 50 ng/ml of macrophage colony-stimulating factor (Immunotools).

To prepare ΔSyk cells we used the Addgene all-in-one CRISPR/Cas9 vector (#79144) as described (Giuliano *et al*, 2019). Briefly, 3 sgRNAs were designed to target mSyk using CRISPick. Control vectors targeting firefly luciferase were obtained from Addgene (# 80248 / 80173 / 80261). Oligos were annealed and phosphorylated before ligation into the plasmid and sequencing. The CRISPR plasmids were then transfected into RAW cells. After 24 h cells were sorted using EGFP fluorescence and an ARIA IIu Becton Dickinson cell sorter into 96-well plates (1 cell / well). Clones grew in 2-4 weeks. They were amplified and checked for Syk expression using Western blots.

### Phagocytosis assays

To assay FcγR-mediated phagocytosis, latex beads (Sigma LB30, 3 µm) were opsonized (10^6^ beads /µl) using 1mg/ ml mouse-IgG (Agrisera AS10 912) for 45 min at 37°C in PBS. Beads were then washed with PBS and resuspended in PBS / BSA (1 mg/ml).

SRBCs were from Eurobio (SB068). They were kept in Alsever solution and used within 3 weeks. For opsonization, 1.10^6^ SRBCs/ µL in opsonisation buffer (50% DMEM /50% PBS / 1 mg BSA/ml) received 1 mg/ml of rabbit anti-SRBCs (MP Biomedicals). After 1h at 37°C, opsonized SRBCs were washed and finally resuspended in opsonisation buffer.

To assay phagocytosis, macrophages were plated on coverslips, washed twice with prewarmed DMEM containing 10 mM Hepes then overlaid with opsonized latex beads (50 /macrophage) or SRBCs (10/ macrophage). Phagocytosis was synchronized by centrifugation for 1 min at 300 g before incubating the plates at 37°C for 5 min (SRBCs) or 15 min (for latex beads). Plates were then placed on ice and washed before labelling extracellular beads using AlexaFluor647 Donkey anti mouse IgG (Jackson Immunoresearch) for 20 min at 4°C (Debaisieux *et al*, 2015). Cells were then washed, fixed with 3.7 % paraformaldehyde and mounted using Vectashield plus (Eurobio). An upright fluorescent microscope (Zeiss Axioimager Z2 with a 63x NA 1.4 objective) was used for counting beads. Images were randomly acquired. At least 100 cells were examined for each condition. The mean number of phagocytosed beads was calculated, and results are expressed as percentage of control cells (mean ± SEM).

To monitor recruitments at the phagocytic cup, cells were transfected with EGFP-Syk or co-transfected with FLAG-Syk and the indicated fluorescent protein (EGFP-Rac1, EGFP-Cdc42, mCherry-Akt or EGFP-PKD- C1ab). After phagocytosis of opsonized SRBCs for 5 min, cells were fixed with 3.7 % paraformaldehyde, permeabilized with 0.2 % saponin, then stained with phalloidin-TRITC before mounting and imaging using a Zeiss LSM880 confocal microscope (with a 63x NA 1.4 objective). Line plots (width: 10 pixels) across the cup and the opposite area of the plasma membrane were generated using Fiji. Enrichment was calculated by dividing the fluorescence peak area at the cup by the fluorescence peak area on the opposite side of the cell. At least 6 phagocyting cells were used to prepare histograms.

### Palmitoylation assays

These experiments were performed essentially as described earlier (Wan *et al*., 2007). Monocyte-derived macrophages or RAW macrophages were allowed to phagocytose SRBCs for 5 min at 37°C as described above. Control cells did not receive SRBCs. Cells were harvested using a cell scraper then lysed on ice with lysis buffer (150 mM NaCl, 1 mM EDTA, 25 mM Hepes, pH 7.2) supplemented with Complete antiprotease (Roche), PhosSTOP (Roche), 1% Triton TX-100, 0.5% CHAPS and 80 nM N-ethylmaleimide (NEM).

Lysates were briefly sonicated (3 x 1 sec), and proteins were precipitated with CHCl3/MeOH and the pellet was resuspended in 4SB (150 mM NaCl, 5 mM EDTA, 50 mM Tris, pH 7.4, supplemented with 4% SDS) containing 50 mM NEM. After 2.5 h at RT on a rotating wheel, samples were subjected to 3 sequential CHCl3/MeOH precipitations to remove NEM. They were split in two. One half was resuspended in buffer H (0.9 M hydroxylamine, pH 7.4, 0.2% TX-100, 0.6 mM HPDP-biotin) and the other half in buffer C (50 mM Tris, pH 7.4, 0.2% TX-100, 0.6 mM HPDP-biotin). After 1 h on the wheel at RT and 2 sequential CHCl3/MeOH precipitations, samples were resuspended in 150 mM NaCl, 1 mM EDTA, 25 mM Hepes, pH 7.2 containing 0.2 TX-100 and 0.05 % SDS, then received streptavidin-agarose (Thermo scientific).

Beads were washed and finally resuspended in 30 µl of reducing SDS-PAGE sample buffer; proteins were separated on 10% acrylamide gels before Western blot. Syk was stained using rabbit polyclonal anti-Syk (CST# 2712) followed by mouse anti-rabbit light chain-peroxidase. Syk phosphorylation on mSyk-Tyr 519/520 was followed using CST# 2710 and mouse anti-rabbit light chain-peroxidase.

### Sulfenylation assays

After SRBC phagocytosis, macrophages were lysed in 1 mM EDTA, 150 mM NaCl, 50 mM Tris, pH 7.5 supplemented with Complete antiprotease (Roche), PhosSTOP (Roche) and 0.5% Triton TX-100. Lysates were treated with 1mM DCP-Bio1 (Hourihan *et al*., 2016) for 1 h at 4°C on a rotating wheel, before anti-Syk immunoprecipitation as described below. The biotin moiety of DCP-Bio1 was stained with Extravidin-Peroxidase. Syk was stained using rabbit polyclonal anti-Syk (CST# 2712) followed by mouse anti-rabbit light chain-peroxidase.

### Syk immunoprecipitation

mSyk and hSyk were immunoprecipitated using a rabbit monoclonal anti Syk (CST#13198) and a mouse monoclonal anti hSyk (4D10), respectively. Lysates were incubated with the antibodies for 1 h at 4°C on the wheel, before adding magnetic beads covered by protein A/G (Pierce 88802) and 1 h at 4°C on the wheel, before washes with lysis buffer, SDS /PAGE and Western blotting.

### Assays for Syk catalytic activity

The FRET assay was performed essentially as described (Xiang *et al*., 2011). ΔSyk RAW cells on coverslips were transfected with the Syk biosensor and the indicated version of FLAG-Syk. After 24h, they were allowed to phagocytose IgG-opsonized SRBCs for 5 min as indicated, then washed and fixed overnight with paraformaldehyde (3.7 %) that was then neutralized with 50 mM ammonium chloride. After washes with PBS, coverslips were mounted in Vectashield plus, then observed using a Zeiss LSM880 confocal microscope fitted with a 63x NA 1.4 objective. Both ECFP and FRET fluorescence were recorded at 475 nm and 530 nm upon excitation at 435 nm. The sensor phosphorylation inhibits the FRET between ECFP and YPET with a concomitant increase in ECFP fluorescence (Xiang *et al*., 2011). Results are expressed as mean ± SEM (n=30-35 cells) of the ratio of fluorescence ECFP / FRET.

### RT-qPCR

RT-qPCR was performed as described earlier (Schatz *et al*, 2023). Briefly, RNA was first extracted using Trizol, then transcribed into DNA using all in one RT Master Mix (Applied Biological Materials). DHHC20 primers for qPCR were designed using primer blast. qPCR was performed using SybrGreen Master Mix as recommended by the manufacturer. Large Ribosomal Subunit Protein EL27 (RPL27) transcripts were used to normalize data. Primers are listed in Table S1.

### Molecular dynamics simulations

The crystal structure of Syk (PDB 4FL3) was used to perform molecular dynamics simulations. First the missing residues were modelled using CharmmGUI (Lee *et al*, 2016) as well as the sulfenylation of the Cys608. Ligands (AMP-PNP, Mg^2+^ and crystal water molecules) present in the PDB structure were preserved for the simulation. Both systems (WT and sulfenylated protein) were immersed in a water box (TIP3P model) surrounding the protein with 20 Å of side edge at a physiological salt concentration ([NaCl] = 0.154 M). Each system was parameterized using the Charmm36m force field (Huang *et al*, 2017) and electrostatic neutrality was achieved by completing with few additional Na+ or Cl− ions depending on the system total charge. Periodic boundary conditions in conjunction with particle mesh Ewald method were set up for each system. After energy minimization (50,000 steps of conjugate gradients) at 0°K and a gradual heating to 310 K, an equilibration phase of 250 ps was achieved. Production runs (used for the analysis) were carried out in the isobaric–isothermal ensemble, at constant temperature (310 K) and pressure (1 atm), using Langevin dynamics and Langevin piston as implemented in NAMD 3.0α12 (Phillips *et al*, 2020) for a total period of 500 ns.

Principal Component Analysis (PCA) based on 3D coordinates from the simulations was achieved using a homemade Python script based of the ProDY package (v2.2.0), allowing for Essential Dynamics Analysis (EDA) (Zhang *et al*, 2021), which was applied to all frames of the production run for both systems by using all CA atoms after superposition of coordinates to the first frame. Root-mean square fluctuations (RMSF) were deduced from the first three normal modes computed by EDA allowing us to evaluate the global mobility of the residues during the simulation. Trajectory analyses and EDA visualization were performed with the Visual Molecular Dynamics program (VMD, v1.9.4, (Humphrey et al, 1996))

## Acknowledgements

This work was funded by the CNRS. We are grateful to Stéphane Bodin, Laurence Abrami and members of the APIR team for discussions and suggestions. The authors declare that they have no conflict of interest.

---

**Supplemental Figure 1.**
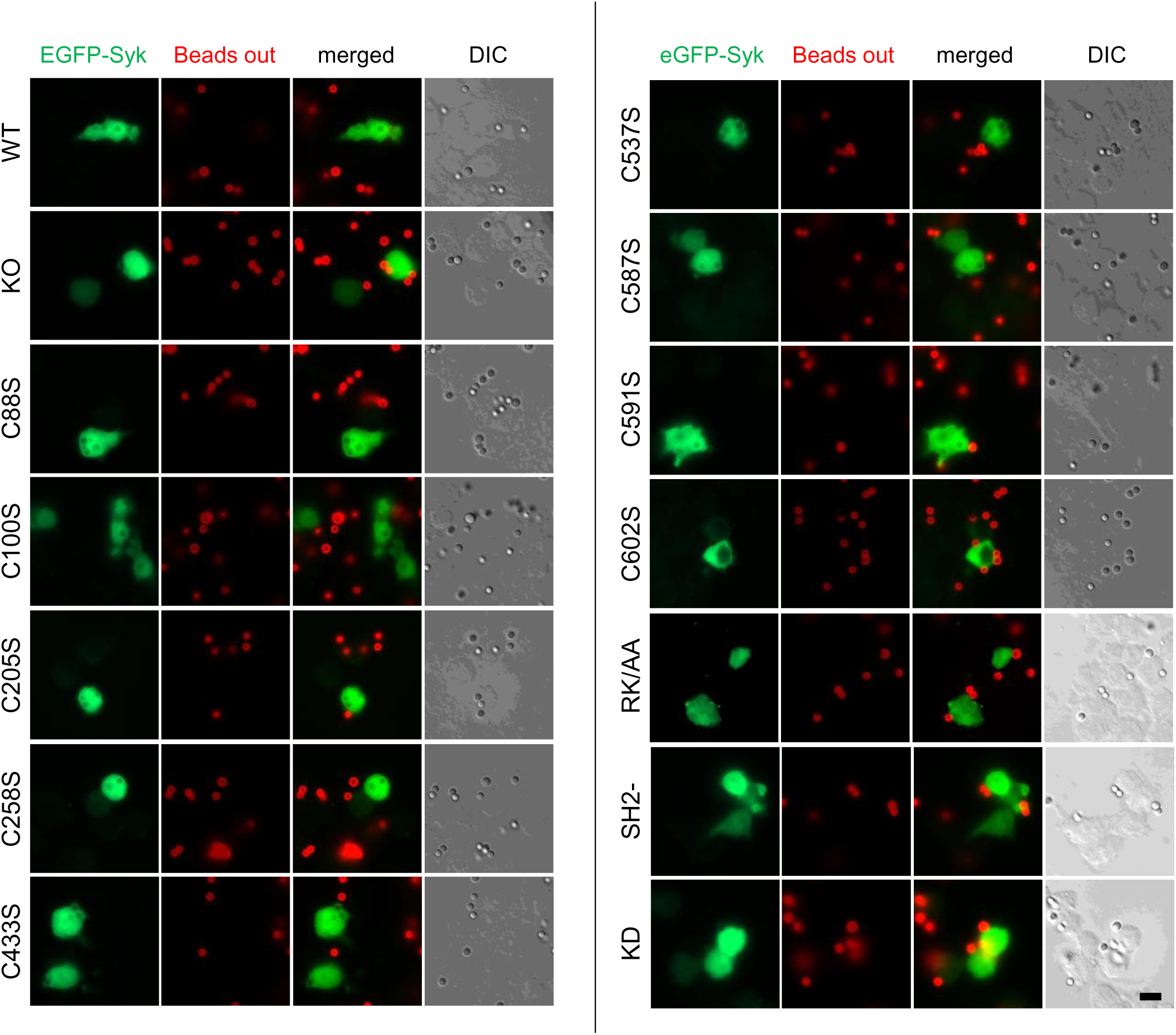
Representative phagocytosis assay. ι1syk RAW 264.7 macrophages were transfected with the indicated version of EGFP-Syk. 18h later, cells were allowed to phagocytose IgG-opsonized 3 µm latex beads for 15 min at 37°C. Extracellular beads were then labelled on ice using Alexafluor647 anti-mouse IgG antibodies (Beads out). Macrophages were then fixed and examined using an upright fluorescent microscope. Total beads were counted using differential interference contrast (DIC) images. Bar, 10 µm.

**Supplemental Figure 2.**
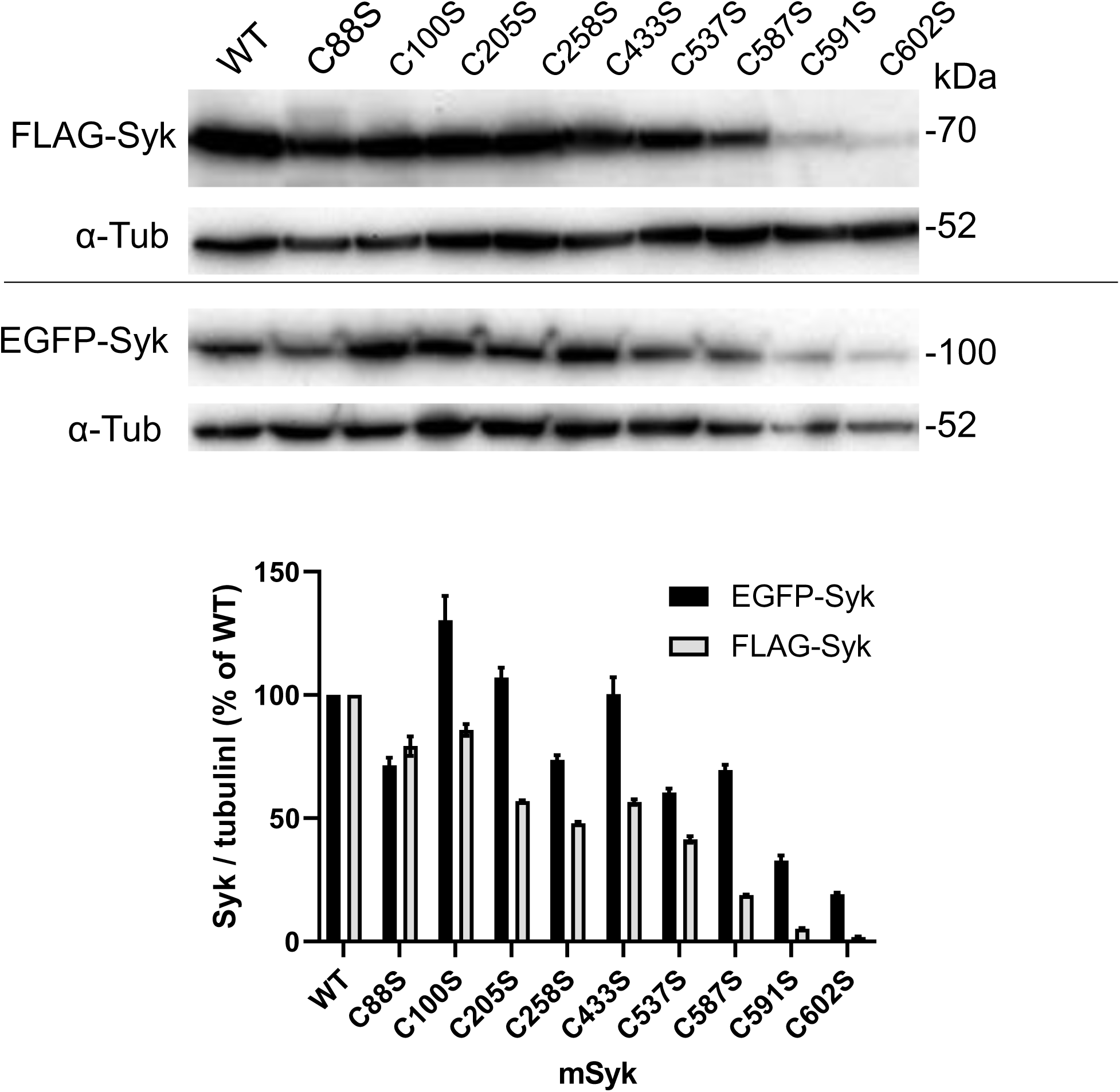
Expression level of Syk-Cys mutants. ι1Syk RAW cells were transfected with the indicated mutant of EGFP-Syk or FLAG-Syk, and cells were harvested for Western blots 24h after transfection. The graph shows the quantification from n=2 blots.

**Supplemental Figure 3a.**
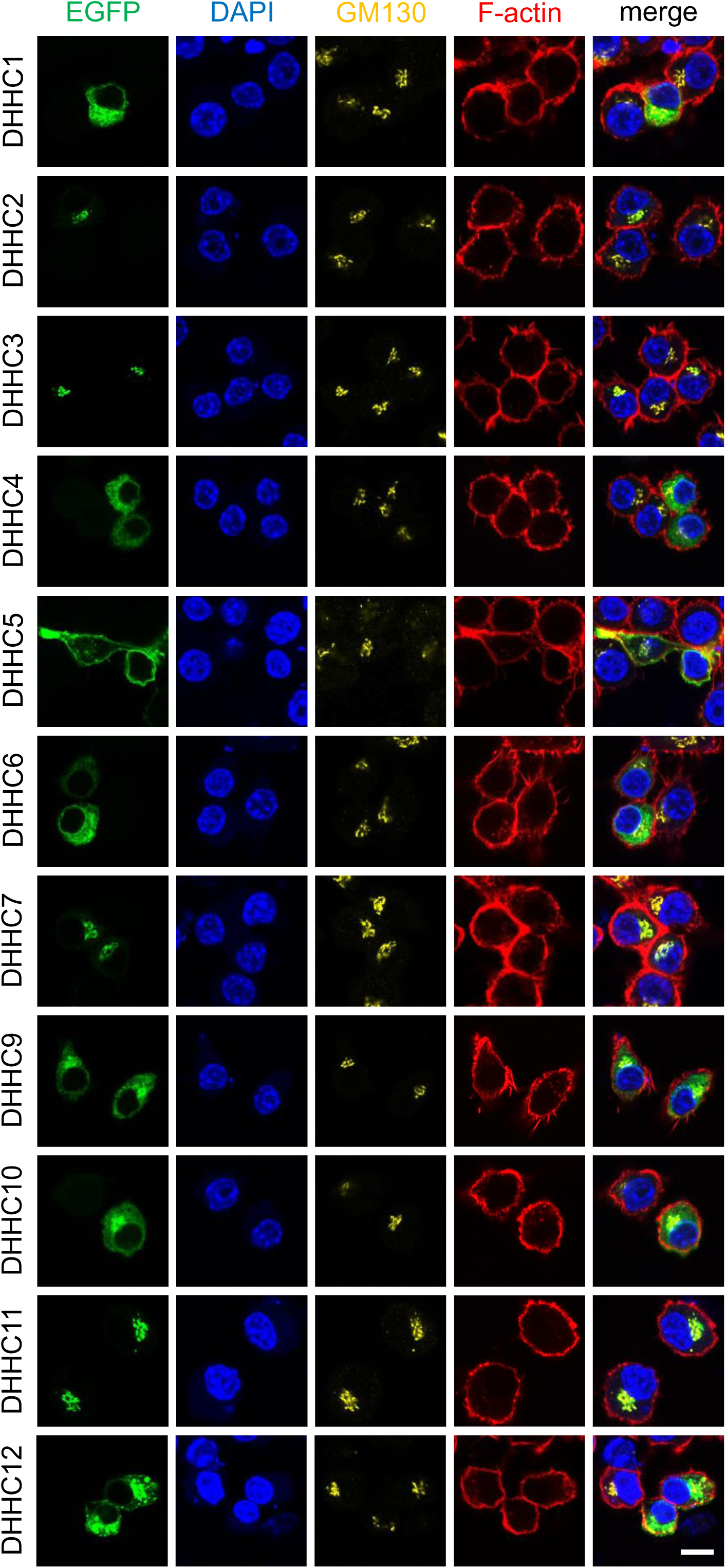
Localization of DHHC enzymes in RAW 264.7 macrophages. Cells were transfected with the indicated EGFP-DHHC. After 24h, cells were labeled with anti-GM130, DAPI and fluorescent phalloidin, then examined by confocal microscopy. Bar, 10 μm.

**Supplemental Figure 3b.**
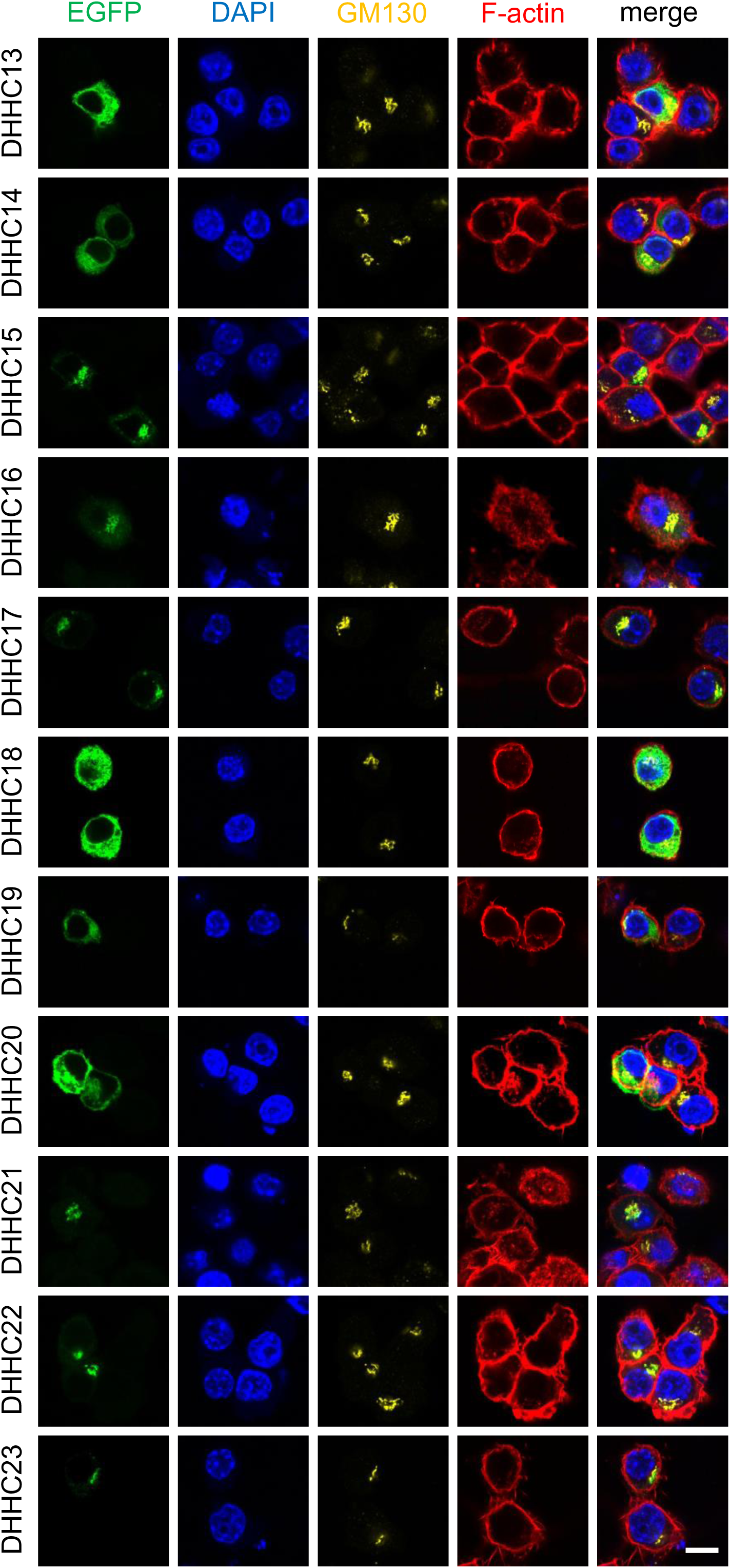
Localization of DHHC enzymes in RAW 264.7 macrophages. Cells were transfected with the indicated EGFP-DHHC. After 24h, cells were labeled with anti-GM130, DAPI and fluorescent phalloidin, then examined by confocal microscopy. Bar, 10 μm.

**Supplemental Figure 4.**
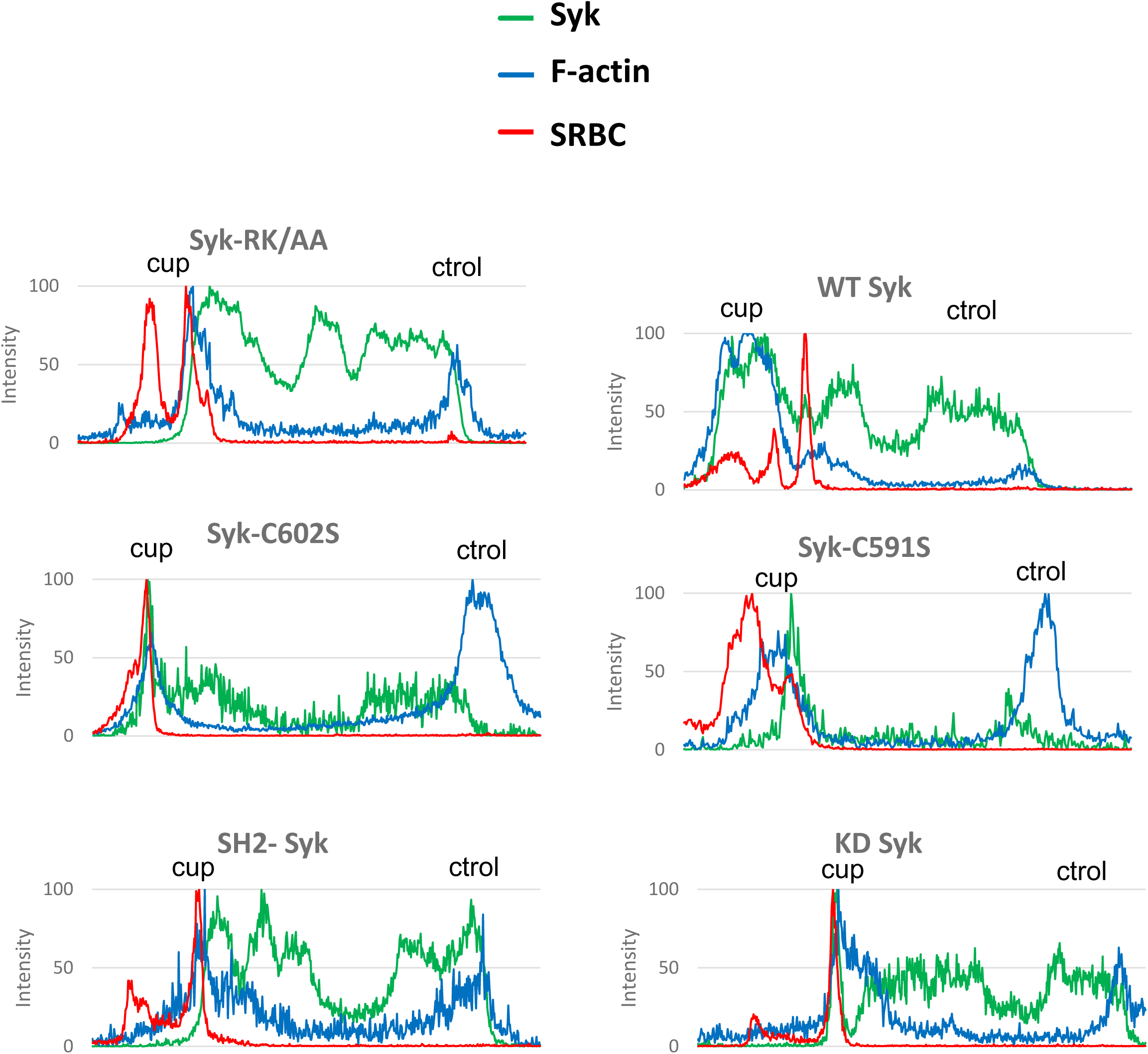
Representative line plots (width: ten pixels) across cells, enabling to monitor Syk, F-actin or SRBC accumulation at the cup (cup) compare to a non-phagocytic region (ctrol) of the plasma membrane (Figure 8).

**Supplemental Figure 5.**
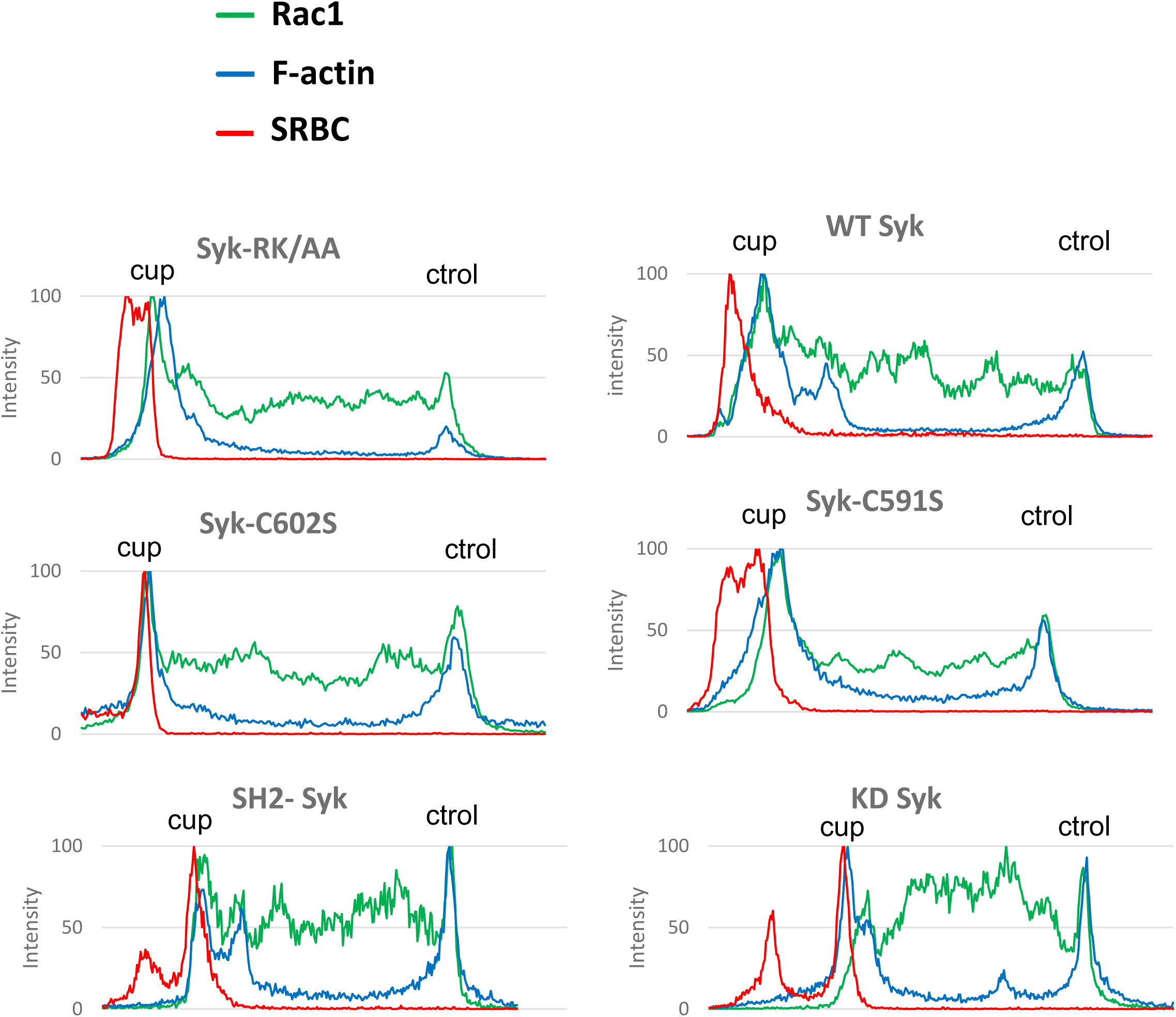
Representative line plots (width: ten pixels) across cells, enabling to monitor Rac1, F-actin or SRBC accumulation at the cup (cup) compare to a non-phagocytic region (ctrol) of the plasma membrane (Figure 9A).

**Supplemental Figure 6.**
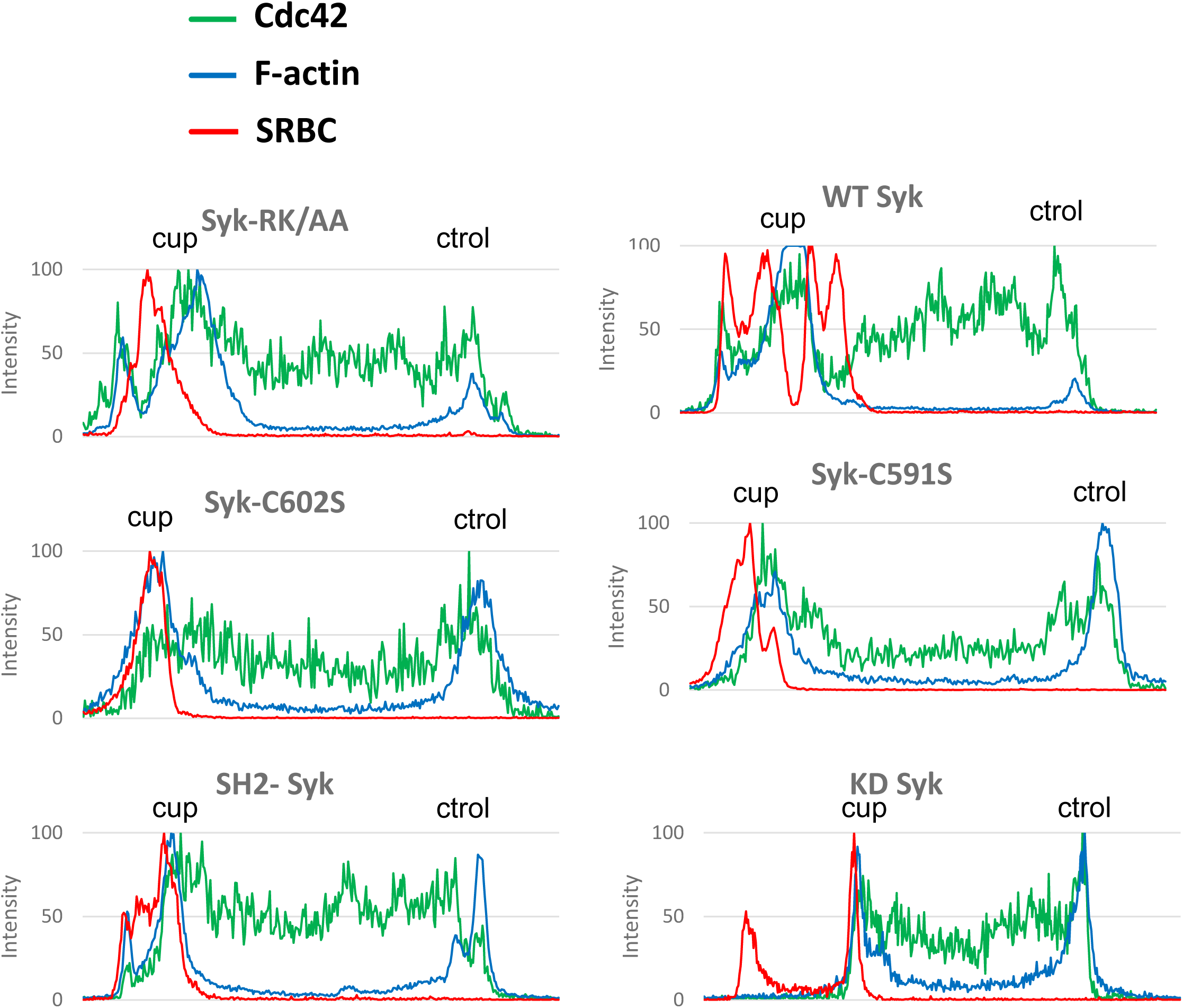
Representative line plots (width: ten pixels) across cells, enabling to monitor Cdc42, F-actin or SRBC accumulation at the cup (cup) compare to a non-phagocytic region (ctrol) of the plasma membrane (Figure 9B).

**Supplemental Figure 7.**
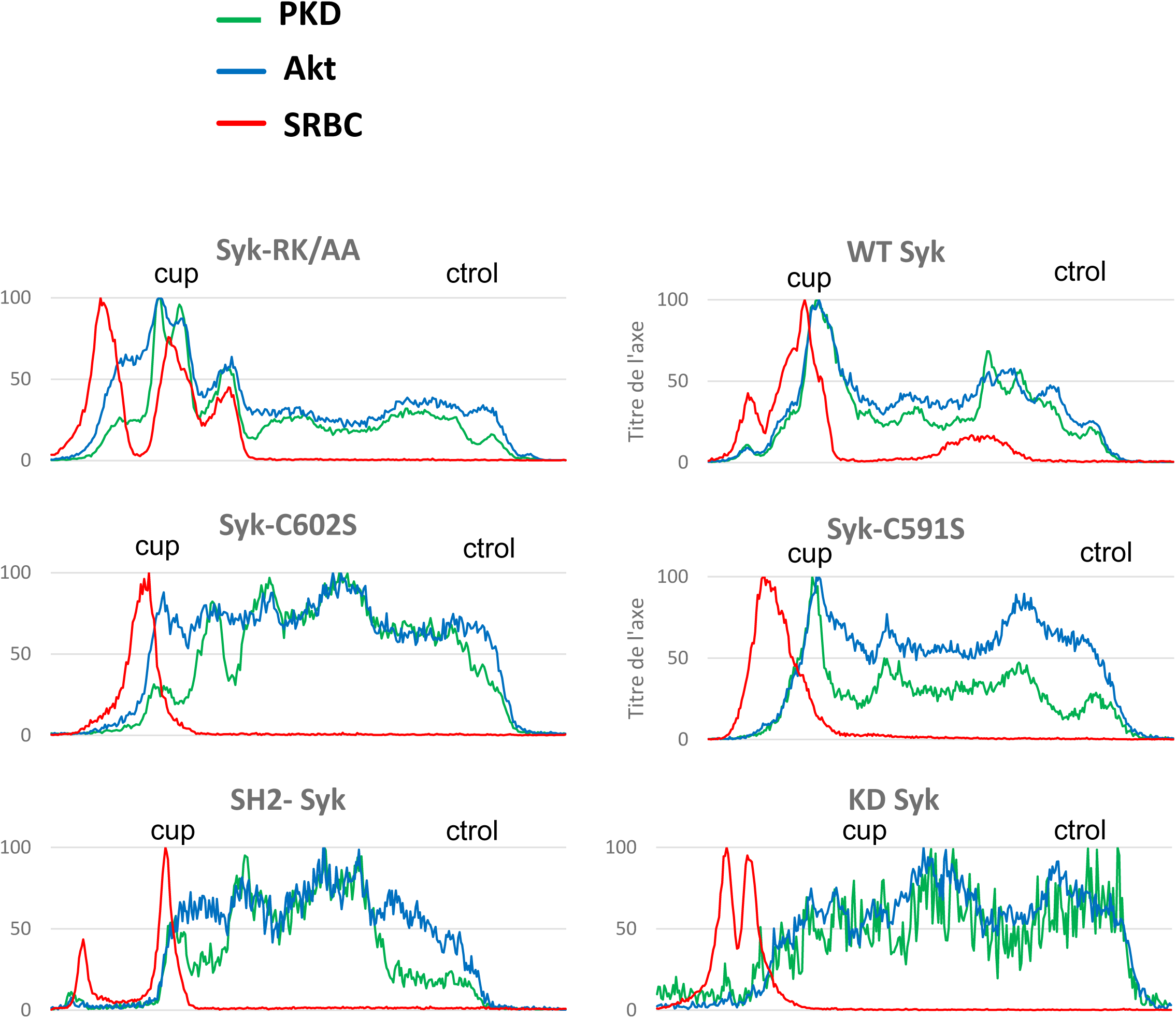
Representative line plots (width: ten pixels) across cells, enabling to monitor PKD, Akt or SRBC accumulation at the cup (cup) compare to a non-phagocytic region (ctrol) of the plasma membrane (Figure 10A).

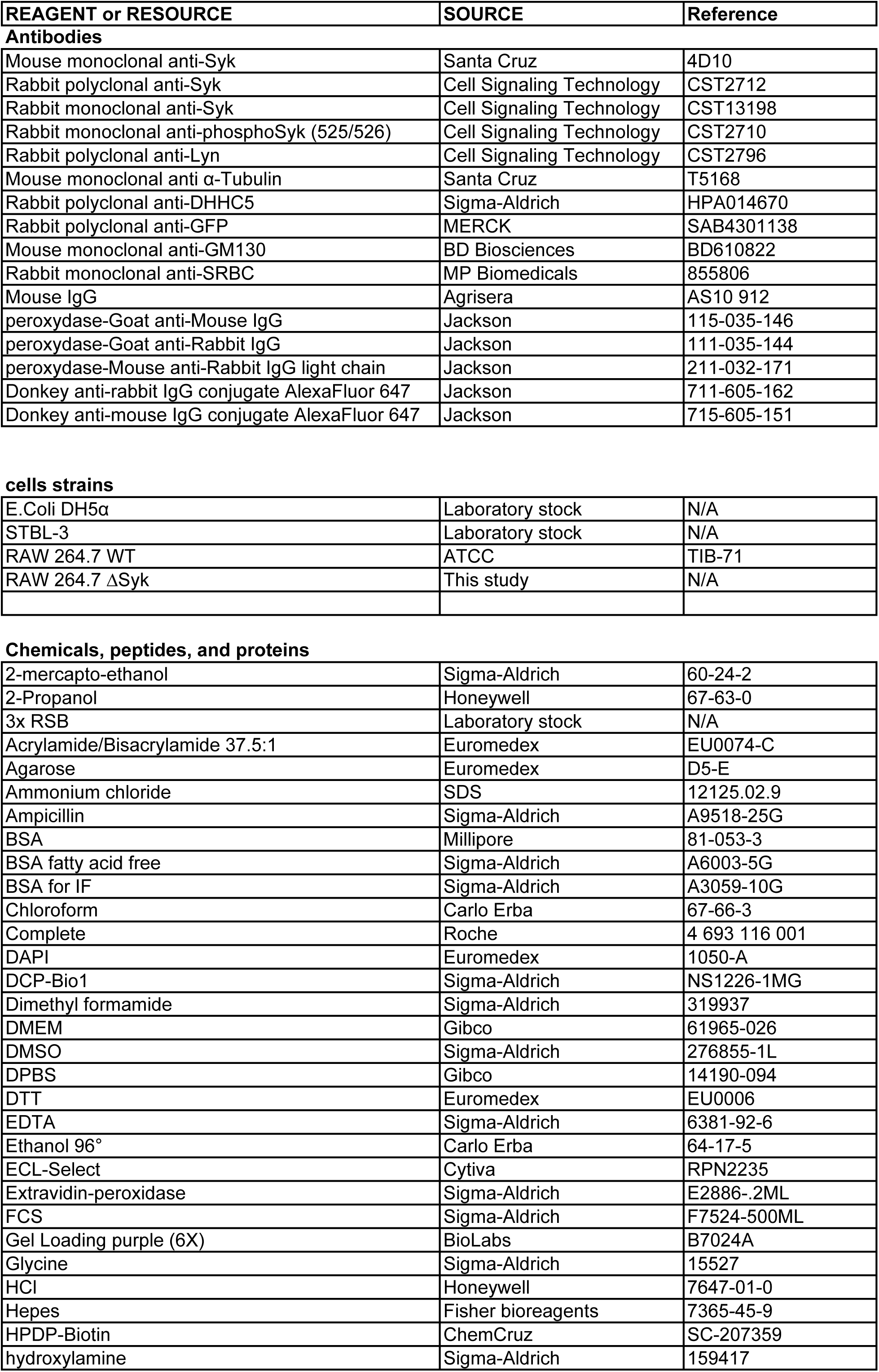
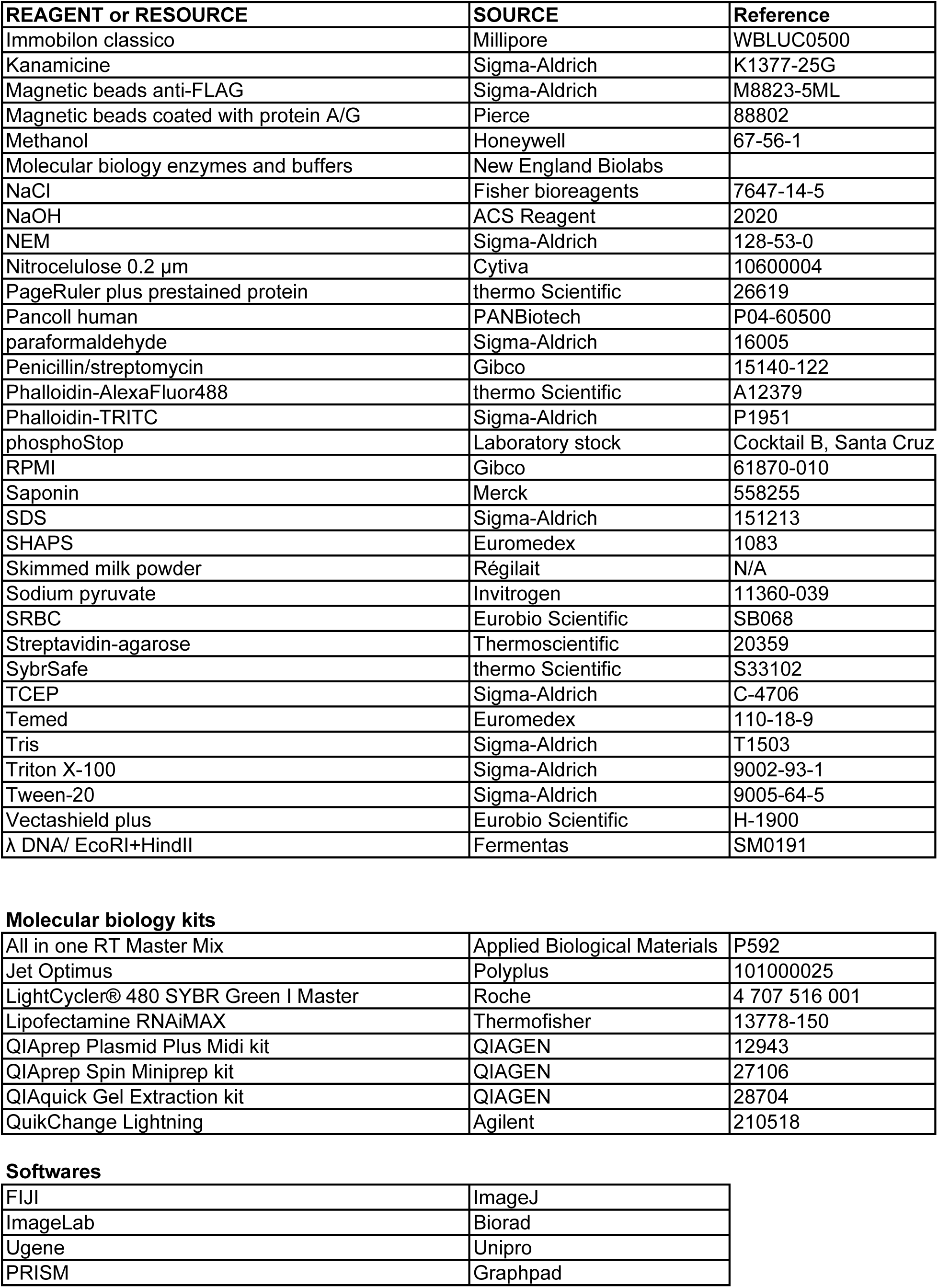
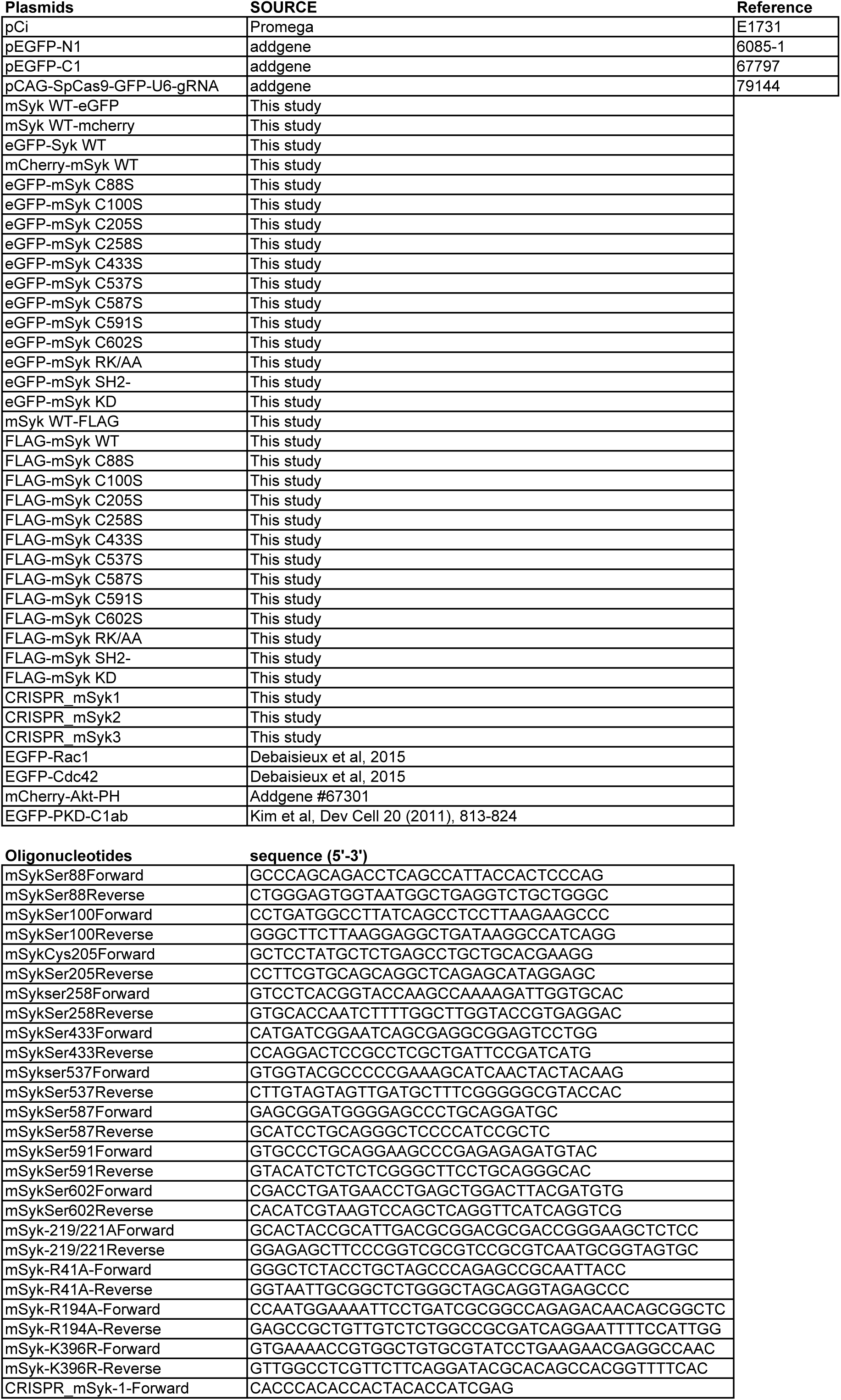
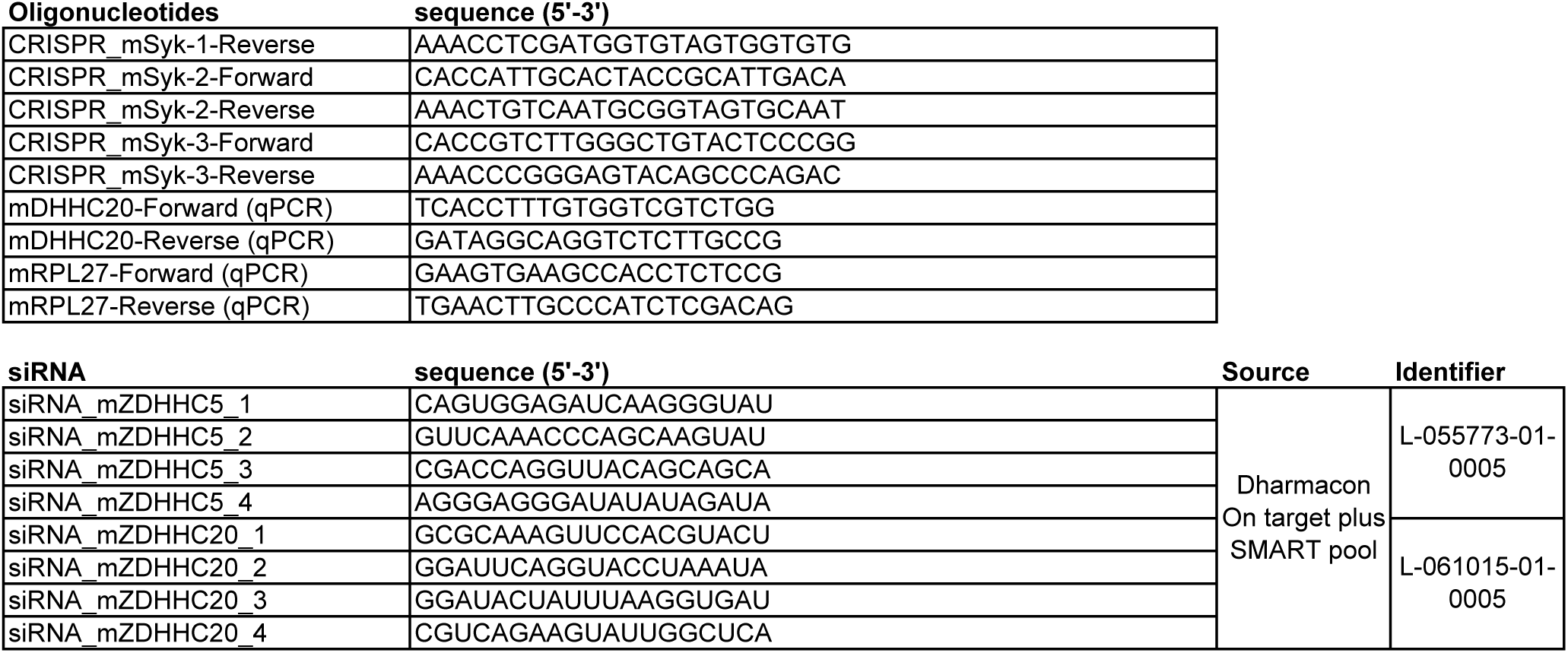

